# Macromolecular Crowding-Affected Mobility of Bcnt/Cfdp1, a Disordered Component of the Srcap Complex, from the Nucleus to Cytosol in Subcellular Fractionation

**DOI:** 10.1101/2025.04.28.651124

**Authors:** Shintaro Iwashita, Takehiro Suzuki, Kouichi Nishimura, Ayaka Onishi, Kyoka Tsukuda, Yoshito Kakihara, Yoshimitsu Kiriyama, Yoshiharu Ohoka, Kentaro Nakashima

**Author notes:** Correspondence &. Department of Pharmacy, Faculty of Pharmacy, Kindai University, Osaka, 577-8502, Japan (Kyoka Tsukuda). These authors contributed equally to the analysis (Shintaro Iwashita, Takehiro Suzuki). Tokushima Bunri University at Sanuki City campus moved to Takamatsu City on 1st April 2025. New address: 8-53 Hamanocho, Takamatsu, Kagawa, 760-0011, Japan.

## Abstract

Metazoan *Bucentaur/Craniofacial Development Protein 1* (*Bcnt/Cfdp1*) is crucial for embryonic development, and its yeast homolog protein, SWC5, functions as part of the SWR1 complex in histone exchange reactions. Bcnt/Cfdp1, an intrinsically disordered protein (IDP) subject to various post-translational modifications, contains a putative nuclear localization sequence. Nevertheless, its subcellular localization remains ambiguous. We investigated its intracellular trafficking using diverse approaches. In C2C12 myoblasts, fluorescent protein-tagged Bcnt/Cfdp1 was predominantly localized within the nucleus, preferentially in regions of low DAPI density in myoblasts, and the localization persisted in differentiated multinucleated myotubes. Conversely, detergent-based biochemical fractionation of C2C12 and HEK293T cells indicated a substantial presence of Bcnt/Cfdp1 in cytoplasmic fractions. Quantitative proteomic analysis of fractionated proteins revealed that six IDPs, including a PEST-containing nuclear protein, demonstrated similar distribution patterns to Bcnt/Cfdp1; despite being annotated as nuclear proteins, they were detected in cytoplasmic fractions. Direct treatment of both cell lines with digitonin enabled the collection of released proteins in three steps to accurately isolate cytoplasmic proteins. Bcnt/Cfdp1 exhibited distinct extraction patterns compared to cytoplasmic soluble protein extracts in Ficoll/sucrose solution, while no differences were noted when cells underwent digitonin treatment in an isotonic solution. Quantitative proteomic analysis of these extracts highlighted the mobility or integrity of each protein within the cytoplasm. Consequently, although Bcnt/Cfdp1 is essential within the nucleus, our findings suggest its additional presence in the cytoplasm. We hypothesize that the elastic properties of Bcnt/Cfdp1 facilitate its translocation from the nucleus during subcellular fractionation, which is affected by macromolecular crowding.

## INTRODUCTION

Organisms have evolved cellular regulatory systems to maintain sophisticated homeostasis in response to environmental stimuli, such as heat and radiation, and stimuli generated by intracellular biosynthesis, such as oxidative stress. Cellular gene-dependent macromolecules dynamically shuttle between the nucleus and cytoplasm during biosynthesis and are recruited to appropriate cellular compartments with high efficiency in response to intracellular and extracellular stimuli. Recent evidence has revealed that these macromolecular behavior dynamics depend on the generated condensates or liquid-liquid phase separation (LLPS), which allows the inclusion and exclusion of necessary and unnecessary components, respectively [1–3].

Among the myriad cellular macromolecules, intrinsically disordered proteins (IDPs) and proteins containing intrinsically disordered regions (IDRs) have attracted attention in the context of condensate formation. IDPs and IDRs are frequently found as scaffolding proteins with multivalent macromolecules and RNA in condensates, such as stress granules (SGs) and processing bodies (PBs) in the cytoplasm and paraspeckles in the nucleus [4]. It is noteworthy that the subtle structure changes in IDRs, such as modifications through phosphorylation/dephosphorylation, have the potential to regulate condensate formation [3]. The low-density condensates have porous and permeable properties, demonstrated in SGs and nuclear speckles [5], where most molecules move freely while the concentrations of related molecules are high [6].

In the context of nuclear-cytoplasmic transport, the structure of the nuclear membrane, particularly the nuclear pore complex (NPC), the only gate with bidirectional potential between the cytoplasm and nucleus, has been intensively studied; IDP and IDR have also been found to play crucial roles in this process [7]. The two types of nuclear transport mediated by NPCs are classified as active and passive transport, depending on the requirement of receptors and energy. The barrier allows ≤30–60 kDa molecules to pass through passive diffusion, showing its entropy-driven nature rather than geometric constriction [8]. Notably, IDPs can diffuse through the NPC without requiring energy or specific transport receptors, while folded protein transport has been revealed to be size-dependent [9].

From yeast to humans, metazoan Bucentaur/Craniofacial development protein 1 (Bcnt/Cfdp1)^‡^ displays a high degree of structural disorder in the N-terminal region and an evolutionarily conserved C-terminal amino acid sequence (BCNT domain) [10]. The yeast ortholog SWC5 is a subunit of the SWR1 (SRCAP in humans) chromatin remodeling complex, SWR1-C (SRCAP-C in humans), while BCNT/CFDP1 has often been treated as an orphan [11, 12]. This is partly due to the failure to capture it as a component of SRCAP-C compared with other subunits [13]. The N-terminal DEY/F motifs of SWC5 engage the canonical histone H2A–H2B [14], whereas the C-terminal BCNT-C domain binds histone H3–H4 and DNA [15] and activates the ATPase of SWR1 [16]. This dual function may contribute to the direction of the torsional force converting the H2A–H2B dimer to H2A.Z–H2B [14–17].

*SWC5* data, based on genetic information, provide clues for inferring the functional frame of Bcnt/Cfdp1 from a causal perspective (https://www.yeastgenome.org/locus/S000000435) [18]. Although *SWR**1*** and ***S****WC**5*** are non-essential genes, they play crucial roles under stressful conditions and have versatile functions. SWC5 protein abundance increases in response to DNA replication stress, and each null strain has decreased or increased resistance to certain chemical compounds, which, along with other factors, may affect their lifespan [18]. In addition, tight control of the histone exchange reaction has been demonstrated to be essential for regulating chromatin dynamics [19].

In contrast, *Bcnt/Cfdp1* is essential for metazoan development, as has been demonstrated for the *Drosophila* homolog *Yeti* [20], zebrafish *cfdp1* [21, 22], and mouse *Cfdp1* [23], [https://www.jax.org/strain/033834], suggesting its critical role in pluripotency and differentiation. The following two lines of evidence for *Bcnt/Cfdp1* as a selected gene suggest that it may have versatile functions in homeostasis, similar to *SWC5*. First, it contributes to the persistence of Chinese hamster ovary cell cultures [24] and resistance to Marek’s disease in chickens [25]. Perturbation of *Bcnt/Cfdp1* in cells or tissues, whether through gain or loss, has been demonstrated to modulate cell migration, cell cycle progression, and apoptosis/anti-apoptosis [20–22, 26]. These phenotypes are preferentially associated with the Wnt/Notch/Akt signaling pathway [27]. Notably, the Wnt signaling pathway is suppressed explicitly in extraluminal mesenchymal cells of the valves of zebrafish Bcnt/Cfdp1-deficient embryos. This defect leads to cardiac dysfunction [22]. Second, *BCNT/CFDP1* has been identified as a prognostic marker for several types of cancers [28, 29]. In addition, Srcap-C has been demonstrated to be involved in a wide range of biological processes, including differentiation [30], DNA damage repair [31], and disease development [32].

We previously reported that *Bcnt/Cfdp1* knockdown at the genomic level in a mouse embryonic stem (ES) cell line, Cfdp1-K1, had a significant impact on the expression of numerous genes, with more than 4% of all genes upregulated or downregulated by more than two-fold [33]. Consistent with this result, RNAi depletion of BCNT/CFDP1 in HeLa cells has been reported to have widespread effects on cellular functions [26]. Of note, this phenomenon may be comparable to the natural genetic variation found in the fission yeast, *swc5*, likely due to the insufficient functionality of the histone exchange reaction [34]. Given that the aforementioned biological phenotypes induced by *Bcnt/Cfdp1* perturbations are preferentially related to the Wnt/Notch/Akt signaling pathway, it seems reasonable to consider that they are indirectly or partially due to perturbations in the Srcap-C transcriptional regulatory framework. This consideration means that while we have gained insight into the function of Bcnt/Cfdp1, we still need more information about its fundamental properties. Assessing the intracellular trafficking of target proteins is essential for understanding fundamental physiological networks and functions, such as cell cycle control [35], and clarifying the causes of diseases [36].

Vertebrate Bcnt/Cfdp1 has typical putative nuclear localization sequences in the N-terminal region, such as ^50^GKKRKAQSIPARKRR^64^ of the human ortholog. However, its subcellular localization remains ambiguous (https://www.proteinatlas.org/ENSG00000153774-CFDP1/subcellular) mainly due to the lack of appropriate antibodies against Bcnt/Cfdp1 that cannot be ignored. Recently, we overcame the issue of off-target bands and assigned the Western blotting signals of mouse Bcnt/Cfdp1 using a Cfdp1-K1 cell extract as a potential negative control [33, 37]. In light of these findings, herein, we investigate the intracellular trafficking of endogenous Bcnt/Cfdp1 in the mouse myoblast cell line C2C12, a well-established model of myocyte differentiation and multinucleated myotube formation, and in HEK293T cells. We also discuss the causes of the conflicting results regarding the subcellular distribution of Bcnt/Cfdp1 between different approaches, considering nuclear export trafficking and macromolecular crowding. Furthermore, we address the Bcnt/Cfdp1 function in the Srcap-C framework.

## RESULTS

### Observation of Bcnt/Cfdp1 in the Nucleus via Cytochemical Approaches

Assessing the subcellular trafficking of the target molecule is essential for clarifying its functional framework, and examining the endogenous ones is imperative. However, no reliable antibodies against Bcnt/Cfdp1 are currently available for immunocytochemistry. Previously, we detected tag-free mouse Bcnt/Cfdp1 primarily in the nuclei of human HEK293T cells by immunocytochemistry with two antibodies generated against two peptides of mouse Bcnt/Cfdp1: one is specific to mice and located in the N-terminal region and the other is common to mice and humans in the C-terminus (Figure 1A, modified from Nakashima *et al*. [37]).

**Figure 1.**
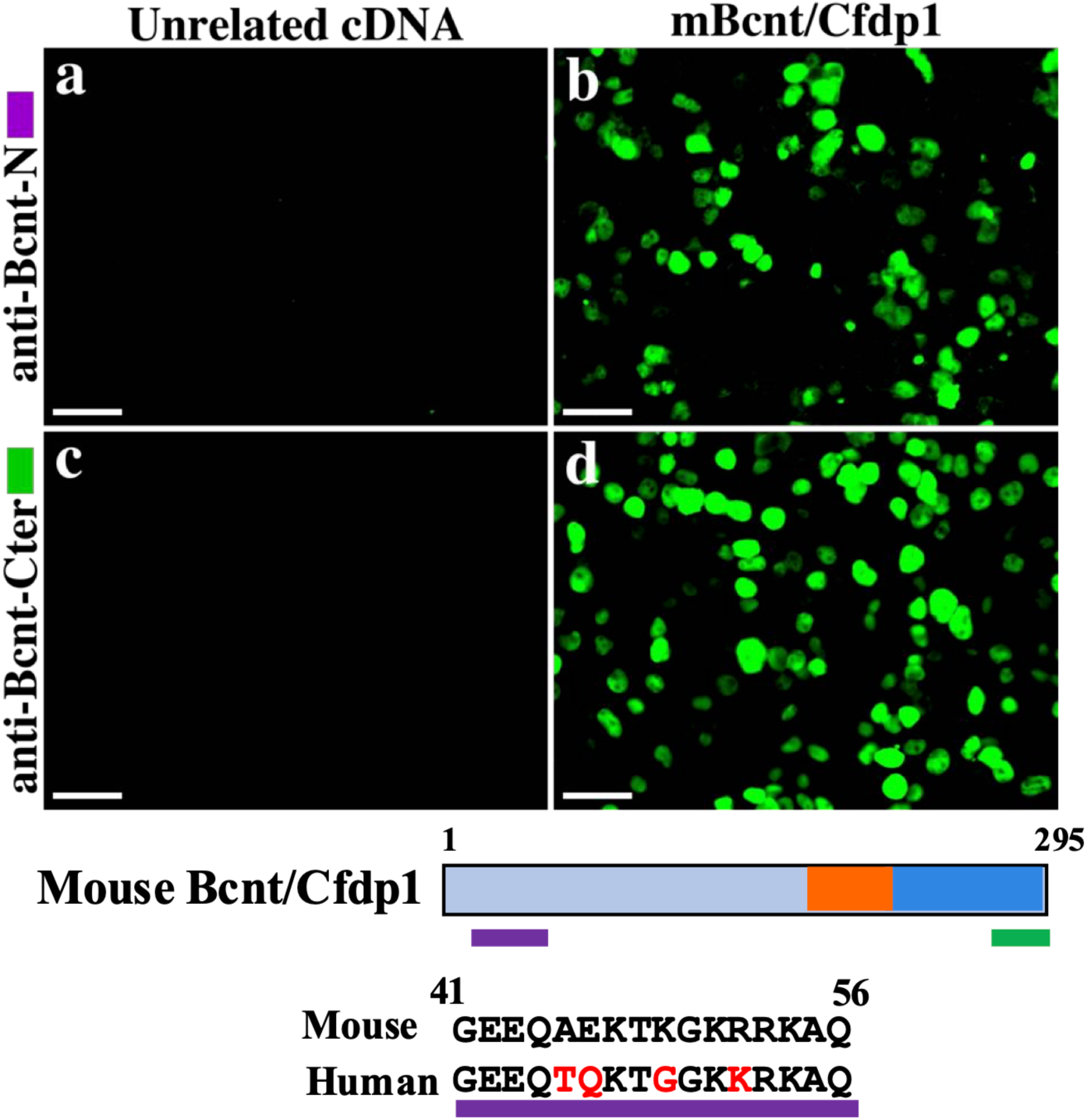
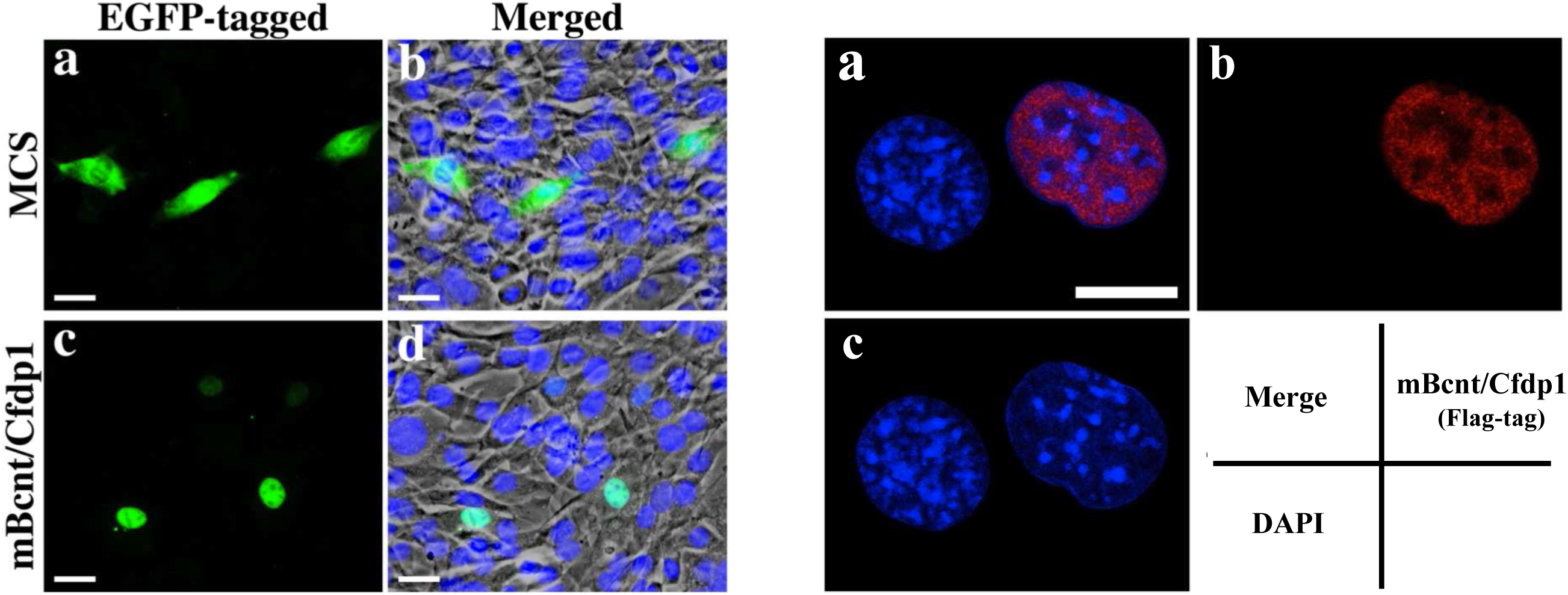
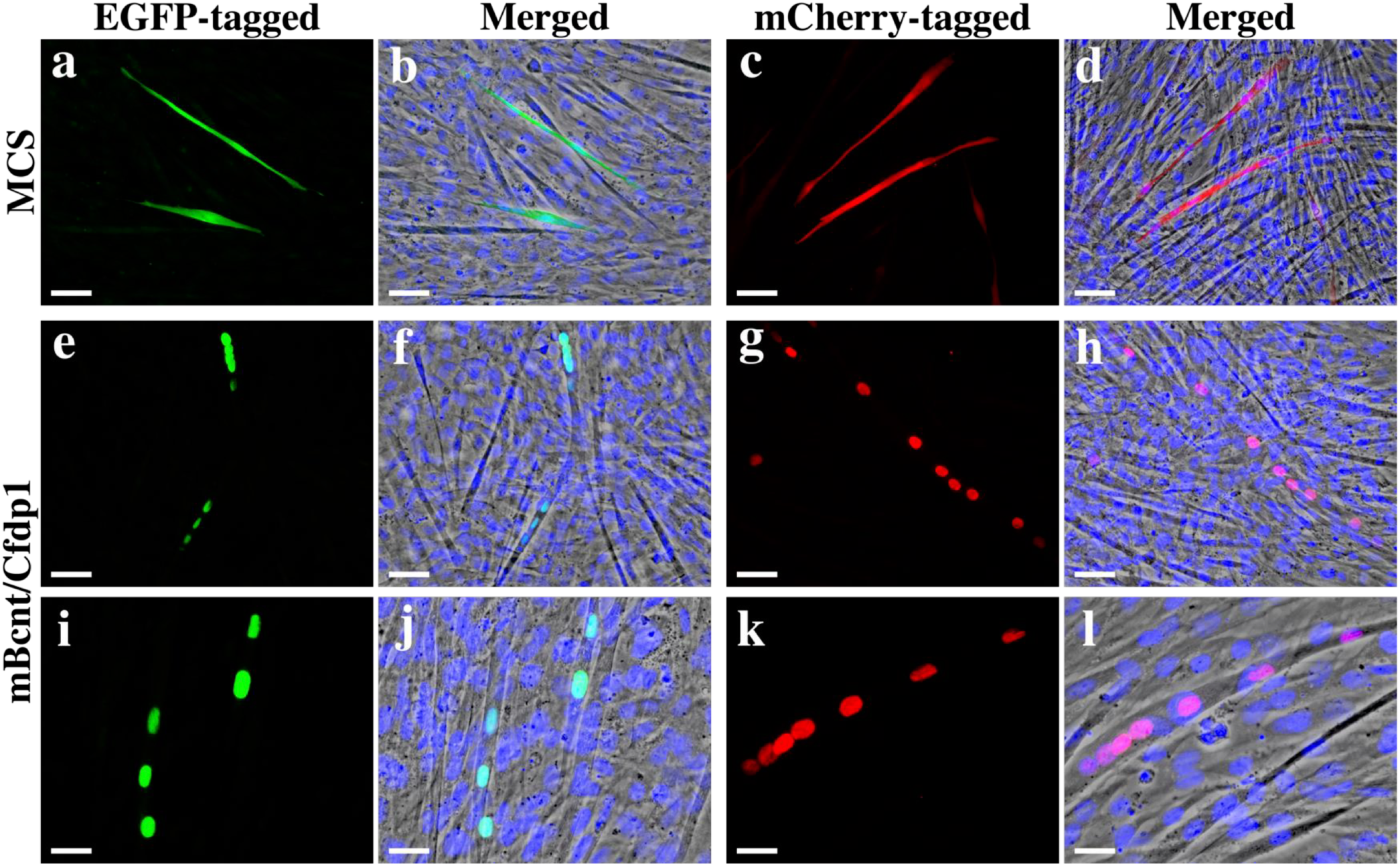
Detection of Bcnt/Cfdp1 primarily in the nucleus by a cytochemical approach. A cytochemical approach was employed to detect the localization of Bcnt/Cfdp1, primarily in the nucleus. **A. Use of tag-free mouse Bcnt/Cfdp1 in HEK293T cells** HEK293T cells were cultured in dishes coated with polylysine, transfected with plasmids encoding tag-free mouse Bcnt/Cfdp1 or unrelated proteins, and immunostained with an anti-Bcnt-N Ab (a, b) or anti-Bcnt-Cter Ab (c, d). The image was visualized using Alexa488-conjugated secondary Ab. The two Abs were generated against two peptides, respectively. The N-terminal amino acid sequence of Bcnt/Cfdp1 was aligned with its human counterpart, with red letters indicating different amino acids. The original images are published by Nakashima et al. [36]. **B. Use of fluorescent protein-tagged mouse Bcnt/Cfdp1 in C2C12 myoblasts** **B-1.** C2C12 cells were cultured in dishes coated with polylysine and transfected with plasmids encoding EGFP-MCS (a, b) or EGFP-Bcnt/Cfdp1 (c, d). Following a 24 h incubation, the cells were fixed with 4% PFA on ice. The samples were immunostained with anti-FLAG Ab followed by CF488A conjugated secondary Ab. The nuclei were stained with DAPI (b, d). The merged images of EGFP and DAPI on phase contrast pictures are shown (b, d). The scale bars indicate 25 μm. **B-2**. C2C12 myoblasts transfected with Flag-mCherry-Bcnt/Cfdp1, similarly cultured and stained as Figure 1B-1, were observed at higher resolution using Leica Mica microscopy. Bcnt/Cfdp1 was stained with anti-Flag Ab followed by Cy3-conjugated secondary Ab (b); the nuclei were stained with DAPI (c), and the merged image of mCherry and DAPI is shown (a). The scale bar indicates 10 μm. **C. In differentiated C2C12 myotubes** After 24 h of culturing the EGFP-MCS (a, b), EGFP-Bcnt/Cfdp1 (e, f, i, j), mCherry-MCS (c, d) or mCherry-Bcnt/Cfdp1 (g, h, k, l) transfectants, the medium was replaced with differentiation medium (DMEM containing 2% horse serum (DM)). The cells were cultured for an additional 4 days with daily DM changes, after which they were fixed with 4% PFA. The samples were immunostained with an anti-FLAG Ab followed by CF488A- or Cy3-conjugated secondary Ab, and the nuclei were stained with DAPI (b, d, f, h, j, l). The merged images of EGFP, mCherry, and DAPI on phase contrast pictures are shown (b, d, f, h, j, l). Scale bars indicate 50 μm (a–h) and 25 μm (i–l).

Here, we used the myoblast C2C12 cell line, a well-known model of myocyte differentiation and multinucleated myotube formation, to analyze the dynamic behavior of Bcnt/Cfdp1 during these processes. We expressed N-terminal FLAG- and fluorescent protein-tagged mouse Bcnt/Cfdp1 (mCherry or enhanced green fluorescent protein (EGFP)) in C2C12 cells. We examined its subcellular distribution using immunostaining with an anti-FLAG antibody. The experimental results showed that, while the control fluorescent multiple cloning site (MCS) was distributed throughout the cell, the fluorescent protein-tagged Bcnt/Cfdp1 was predominantly localized in the nuclei (Figure 1B-1).

The density of the nuclear interior is recognized as a porous medium composed of the chromatin network [6]. The chromatin density is heterogeneous; the inactive part, heterochromatin, is denser, whereas the active part, euchromatin, is sparser. Using high-resolution imaging, we observed mCherry Bcnt/Cfdp1 and DAPI in the nucleus of C2C12 myoblasts and found their segregation pattern; mCherry Bcnt/Cfdp1 was present in the area with weaker DAPI signals (Figure 1B-2).

**Figure 2.**
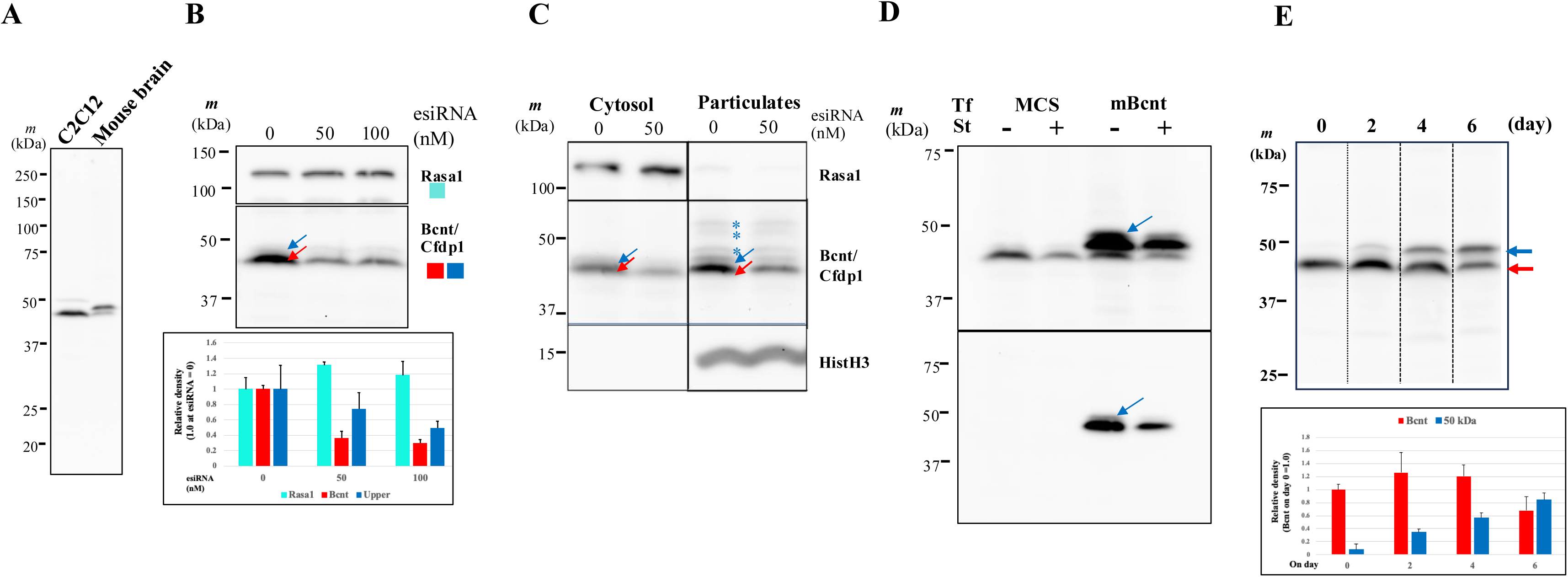
Validity of Western blotting monitoring in C2C12 cell subcellular fractionation. **A. Western blotting signals of Bcnt/Cfdp1 in C2C12 cells** Extracts from whole C2C12 cells (equivalent to 7.5 x 10^4^) or mouse embryo brains (15 μg) were subjected to Western blotting analysis using A305-624A-M as an anti-Bcnt/Cfdp1 Ab. **B. Effects of esiRNA knockdown of Bcnt/Cfdp1 in C2C12 cells.** C2C12 cells were treated with 0, 50, or 100 nM esiRNA for 24 h. Each extract was prepared by scraping off the cells in LS buffer and boiling. After centrifugation, equal amounts of protein (20 μg) were subjected to Western blotting analysis. The upper and middle sections of the image show Western blotting signals using Abs against Rasa1 and Bcnt/Cfdp1, respectively. The red and blue arrows correspond to the target and upper band. At the bottom, densitometric analysis of blots of Bcnt/Cfdp1 (red bar), “upper band(s)” (dark blue), and Rasa1 (light blue) are shown as bar graphics with the mean ± SEM of duplicate experiments. **C. Effects of esiRNA on the cytosolic and particulate proteins** C2C12 cells were cultured in 35 mm dishes and treated with esiRNA (0 or 50 nM). Cytosolic and particulate proteins were obtained using an HFS solution containing 250 μg/ml digitonin, as detailed later in Figures 6-8. Briefly, digitonin-mediated released fluid was the cytosolic fraction, and the particulates were obtained by scrapping off the remaining cell layer in HFS solution followed by centrifugation. The cytosolic fraction was concentrated with TCA precipitation followed by resuspending in LS buffer, while the particulate fraction was directly resolved in LS buffer. Equal relative amounts of each fraction (approximately 15 or 25 μg of protein, respectively) were subjected to Western blotting analysis to assess the degree of separation using anti-histone H3 Ab in addition to that in Fig. 3B. The red and blue arrows correspond to the target and “upper band,” respectively. Asterisks (*) indicate off-target bands. **D. The presence of the “upper band” in the FLAG-tagged Bcnt/Cfdp1** C2C12 cells were transfected with F-MCS or F-Bcnt/Cfdp1 (Tf: transfection). After 24 h of culture, one sample from each group was replaced with serum-free medium (St: serum-starved) and cultured for 6 h. All cell cultures were harvested and subjected to Western blotting analysis using anti-Bcnt/Cfdp1 (upper panel) or anti-FLAG (lower panel) Abs. The blue arrows indicate the “upper band” of F-Bcnt/Cfdp1. **E. The appearance of a cross-reacted 50 kDa band during C2C12 differentiation** C2C12 cell cultures were switched from regular medium to DMEM containing 2% horse serum, designated as day 0. Cell extracts were collected and subjected to Western blotting analysis the following day. The red and blue arrows indicate Bcnt/Cfdp1 and an unrelated 50 kDa band. At the bottom, densitometric analysis of blots of Bcnt/Cfdp1 (red bar) and 50 kDa band (dark blue) are shown as bar graphics with the mean ± SEM of duplicate experiments.

To gain insights into the intranuclear dynamics of Bcnt/Cfdp1, we examined its potential involvement in forming the condensates in the nucleus using the N-terminal BCNT/CFDP1 IDR. Nevertheless, the results were in vain (Supplementary Figure 1S, which will be discussed later). Furthermore, the fluorescent protein-tagged Bcnt/Cfdp1 remained stable during prolonged culture, and multiple signals were visible in the nuclei (Figure 1C). These results indicated that fluorescent protein-tagged Bcnt/Cfdp1 was detected primarily in the nuclei of C2C12 myoblasts and differentiated multinucleated myotubes.

### Validity/Invalidity of Western Blot Monitoring in C2C12 Cell Subcellular Fractionation

To determine the subcellular distribution of endogenous Bcnt/Cfdp1 in cells, we investigated Bcnt/Cfdp1 through the biochemical fractionation of C2C12 and HEK 293T cells, monitored by Western blot analysis, followed by mass spectrometry. Our previous study demonstrated that the Western blotting signal of Bcnt/Cfdp1 corresponds to a doublet of ∼43 kDa (previously described ∼45 kDa) bands in mouse ES cell lines Cfdp1-K1 and mouse brain extracts using A305-624A-M as an anti-CFDP1 antibody [37]. We confirmed that the ∼43 kDa signal band represents Bcnt/Cfdp1 by enriching the immunoreactive band from Cfdp1-K1 cells using two-column chromatography followed by mass spectrometry (Supplementary Figure S2 and Supplementary Table S1). Next, we validated the Western blot signal(s) of C2C12 myoblast extracts obtained with A305-624A-M (Figure 2A-D). The treatment of an endonuclease-derived siRNA (esiRNA) against mouse *Bcnt*/*Cfdp1* significantly decreased the signal of the ∼43 kDa band, but the effects on the lower-mobility band(s) were lesser (Figure 2B and 2C). To confirm whether the lower-mobility band is derived from Bcnt/Cfdp1, which was previously referred to as the “upper band” corresponding to its phosphorylated form [10, 33], we expressed and isolated FLAG-tagged mouse Bcnt/Cfdp1 in C2C12 cells (Figure 2D and Supplementary Figure S3). A small upper band appeared in C2C12 cells expressing FLAG-tagged Bcnt/Cfdp1, but its phosphorylated peptide at serine 245 in the ∼43 kDa band [33] was not captured (data not shown). These results conclude that the upper band exists, but its signal region contains both Bcnt/Cfdp1-derived molecule and off-target(s), which will be discussed later.

Compared to the cytoplasmic soluble fraction, the particulate fraction exhibited several off-targets unaffected by esiRNA treatment (Figure 2C). In addition, during myotube differentiation, the distinct 50 kDa signal gradually increased, whereas Bcnt/Cfdp1 levels decreased in the later stages (Figure 2E and Supplementary Figure S4). These results indicate that Bcnt/Cfdp1 can be effectively monitored in the cytoplasmic soluble fraction of C2C12 myoblasts by Western blotting. However, tracking the target in the nuclear fraction and differentiated cell extracts, including the upper band(s), may not be feasible. Consequently, we focused on the cytoplasmic soluble fraction.

### Distribution of Bcnt/Cfdp1 by Subcellular Fractionation

We performed biochemical fractionation using a widely utilized kit from Thermo Scientific, which may include different detergents in each step [38]. C2C12 or HEK293T cell suspensions were serially fractionated according to the manufacturer’s protocol, with additional washes at each step. The five resulting fractions were subjected to Western blotting analysis using antibodies against Rasa1 (p120 Ras GTPase activating protein) and Gapdh (glyceraldehyde-3-phosphate dehydrogenase) as markers of the cytoplasmic extract, Vdac1 (voltage-dependent anion channel 1) for the membrane extract, NonO (non-POU domain-containing octamer-bound p54nrb) for the soluble nuclear extract, and HstH3 (histone H3) as a marker of the chromatin-bound fraction. Nestin, a linker protein that interacts with the cytoskeleton, was also used and detected in the cytoskeletal fraction of C2C12 cells and the cytoplasmic fraction of HEK293T cells (Figure 3). Under these fractionation conditions, a substantial amount of the ∼43 kDa band was detected in the cytoplasmic fraction, while the nuclear fraction contained other bands (s) of ∼50 kDa (Figure 4A). We subjected these five fractions of C2C12 myoblasts to quantitative mass spectrometry to evaluate the fractionated preparations quantitatively. A total of 4,487 proteins were identified, of which 3,280 were eligible to evaluate relative distribution (Supplementary Table S2). These included five biomarkers (Figure 4B), heterogeneous nuclear ribonucleoproteins, and three paraspeckle proteins (Figure 4C). In sequential fractionation, the components from the previous fractions were easily carried over into the following fraction. Nevertheless, a significant amount of Bcnt/Cfdp1 was extracted in the first fraction, the cytoplasmic fraction (Figures 4A and 4B). These results indicated that Bcnt/Cfdp1 is primarily present in the cytoplasmic fraction of C2C12 cells using an available biochemical subcellular fractionation kit and suggested that this is also the case for HEK293T cells (Figure 3, right panel).

**Figure 3.**
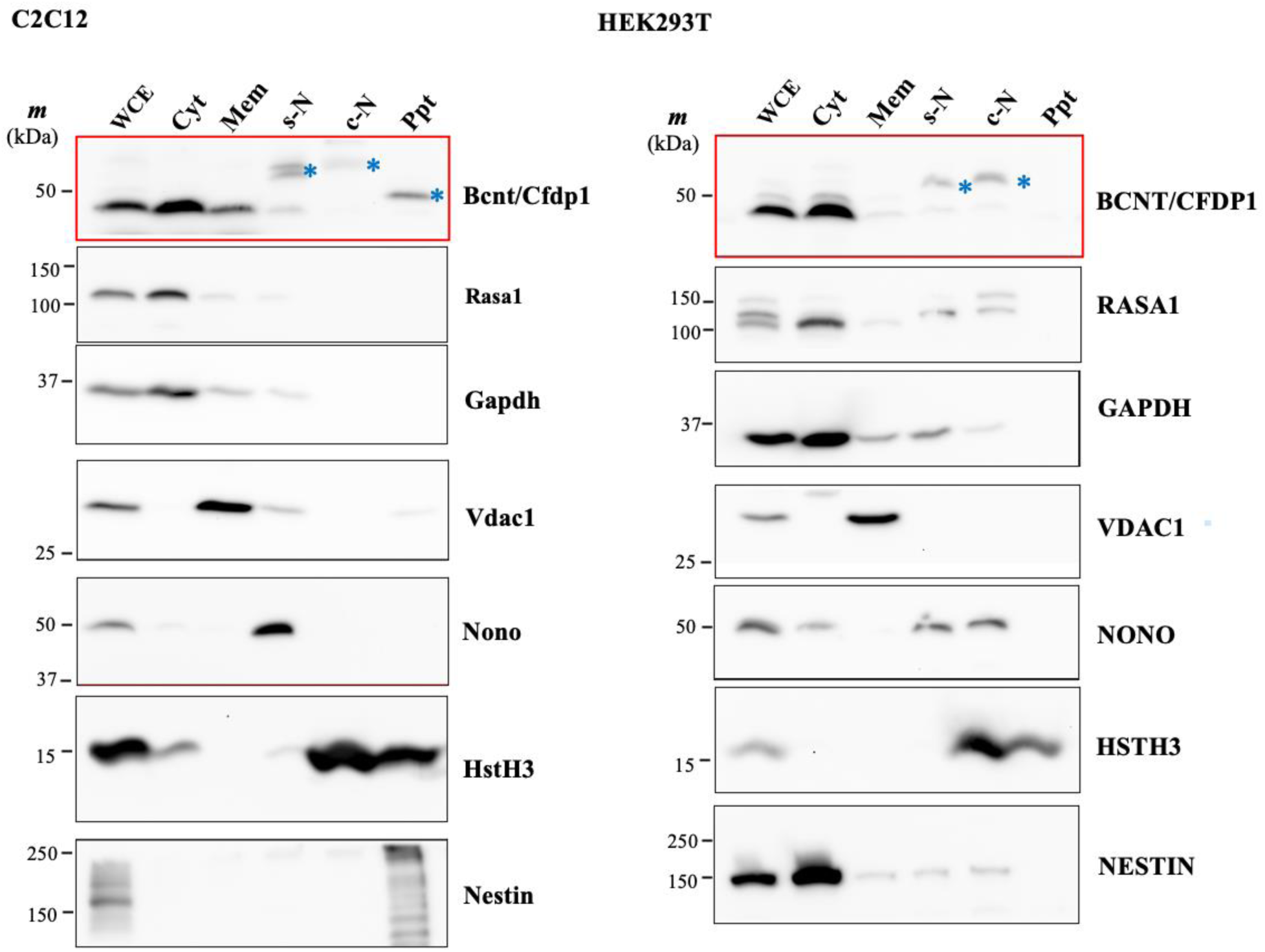
The presence of Bcnt/Cfdp1 in the cytoplasmic fraction determined using a widely used kit. For subcellular fractionation of C2C12 or HEK293T cells, each cell suspension (1 × 10^6^ cells) was subjected to fractionation according to the manufacturer’s protocol, which was slightly modified with additional washes to minimize contamination at each step. Each fraction was concentrated by TCA precipitation and resolved in LS buffer. To show the relative occupancy of all fractions except WCE, equal volumes of each were subjected to Western blotting analysis. The following proteins were accessed: Rasa1 (Ras GTPase activating protein 1), Gapdh (glyceraldehyde-3-phosphate dehydrogenase), Vdac1 (voltage-gated anion channel 1), non-POU domain-containing octamer-binding protein (NonO), HstH3 (histone H3), and Bcnt/Cfdp1 using A305-624A-M. The asterisks in the signals detected with the anti-Bcnt/Cfdp1 Ab indicate the off-target band. Abbreviations at the top of the figure. WCE: whole cell extract, Cyt: cytoplasmic extract, Mem: membrane extract, s-N: soluble nuclear extract, c-N: chromatin-bound nuclear extract, and Ppt: pellet extract.

**Figure 4.**
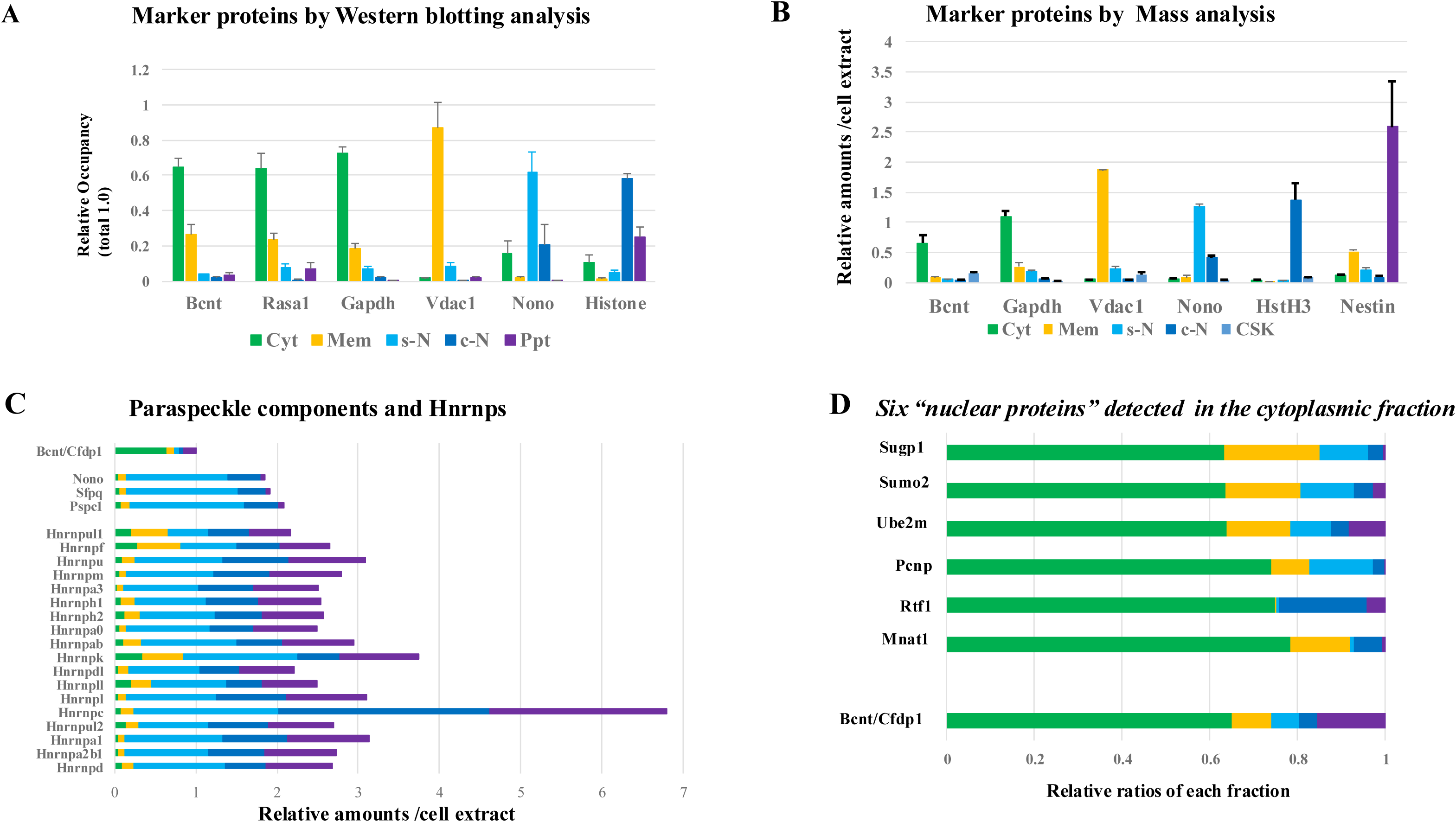
Quantitative analysis of the subcellular fractionated samples accessed by Western blotting or by mass spectrometry. **A.** Distribution of six marker proteins in each fraction accessed by Western blotting Densitometric analysis of Western blots of six biomarker proteins (Bcnt/Cfdp1, Rasa 1, Gapdh, Vdac1, NonO, and HstH3) in C2C12 subcellular fractions of Figure 3 is shown. Data represent the relative occupancy of each marker protein in the five fractions, in which total amounts are set as 1.0, and the bars represent the mean ± SEM of results obtained from five independent experiments. Cyt, cytoplasmic fraction; Mem, membrane fraction; s-N, soluble nuclear fraction; c-N, chromatin-bound fraction; Ppt, precipitates. **B–D** Two independent C2C12 myoblast cultures were fractionated in a manner analogous to that described in Figure 4. Each fraction was subjected to quantitative mass spectrometry analysis. A total of 4,489 proteins were identified, and the average abundance of each component was calculated and displayed graphically. Cyt, cytoplasmic fraction; Mem, membrane fraction; s-N, soluble nuclear fraction; c-N, chromatin-bound fraction; CSK, cytoskeleton. **B.** Distribution of six marker proteins in each fraction accessed by mass analysis Data represent the relative amounts of each marker protein per cell extract in the five fractions, in which total amounts of Bcnt/Cfdp1 are set at 1.0, and bars represent the mean ± SEM of results obtained from two independent experiments. **C.** The intracellular distributions of three paraspeckle components and 18 heterogeneous nuclear ribonucleoproteins The data are shown in relative quantitative amounts, with the total Bcnt/Cfdp1 set at 1.0. The data represent the means of two experiments. **D.** The intracellular distribution of six proteins The intracellular distribution of proteins screened as those detected in the cytoplasmic fraction despite being annotated localized in the nucleus is shown. The relative distribution of each protein is shown from the mean value of the two experiments.

However, this result was in contrast to the above finding that the Bcnt/Cfdp1 was predominantly observed in the nucleus using fluorescent protein labeling and immunocytochemical techniques (Figures 1A–C). To gain insight into this intracellular Bcnt/Cfdp1 mobility during the subcellular fractionation processes in C2C12 cells, we searched for proteins that showed similar distribution to Bcnt/Cfdp1; the proteins annotated or reported to be present predominantly in the nucleus are primarily detected in the cytoplasmic fraction. Of the 3,280 proteins available for quantitative evaluation, 472 showed high relative occupancy (>0.6) in the cytoplasmic fraction, of which six proteins were annotated as present in the nucleus (Figure 4D). They were IDR-containing proteins, such as PEST proteolytic signal-containing nuclear protein, Pcnp [39]. Based on further curation using the integrated cellular localization prediction software BUSCA [40] and experimental evidence from human ortholog data [41], these proteins may localize to the cytoplasm (Supplementary Table S3).

### Intracellular Behavior of Bcnt/Cfdp1 in Digitonin-Mediated Release

To further investigate the cytoplasmic distribution of Bcnt/Cfdp1 strictly, C2C12 and HEK293T cell cultures on dishes were directly treated with digitonin to obtain the cytoplasmic soluble proteins. Digitonin is a unique detergent that selectively binds to the cholesterol in membranes, creating pores in cholesterol-rich membranes that allow macromolecules to pass through [42]. In addition, it has been observed that digitonin does not flip across membranes with low cholesterol content, such as the nuclear envelope [43]. Therefore, digitonin is currently one of the most appropriate methods for isolating the cytosolic proteins of mammalian cells while exerting minimal effects on the intracellular behavior of other proteins, provided that the treatment conditions are appropriately determined.

#### A) In an isotonic solution

C2C12 myoblasts on the dish were exposed to isotonic solutions containing different concentrations of digitonin. The released eluate and corresponding residual cell contents were collected and subjected to Western blotting analysis. A prominent band of ∼43 kDa was observed in the eluate at a digitonin concentration of 25 μg/mL, showing a similar pattern to that of the cytoplasmic soluble proteins Rasa1 and Gapdh (Figure 5A). The ∼43 kDa band was excised from the gel and subjected to mass spectrometry analysis (Figure 5B). The data showed that Bcnt/Cfdp1 was ranked high in the Sum Pep Score of 124 identified proteins annotated as cytoplasmic proteins except for nuclear protein Trip13 (Supplementary Table S4 and Figure 5C). However, the human ortholog TRIP13 has been reported to be distributed diffusely in the cytoplasm, in addition to the nucleus [44]. Thus, these results support the cytoplasmic distribution of Bcnt/Cfdp1 in C2C12 myoblasts.

**Figure 5.**
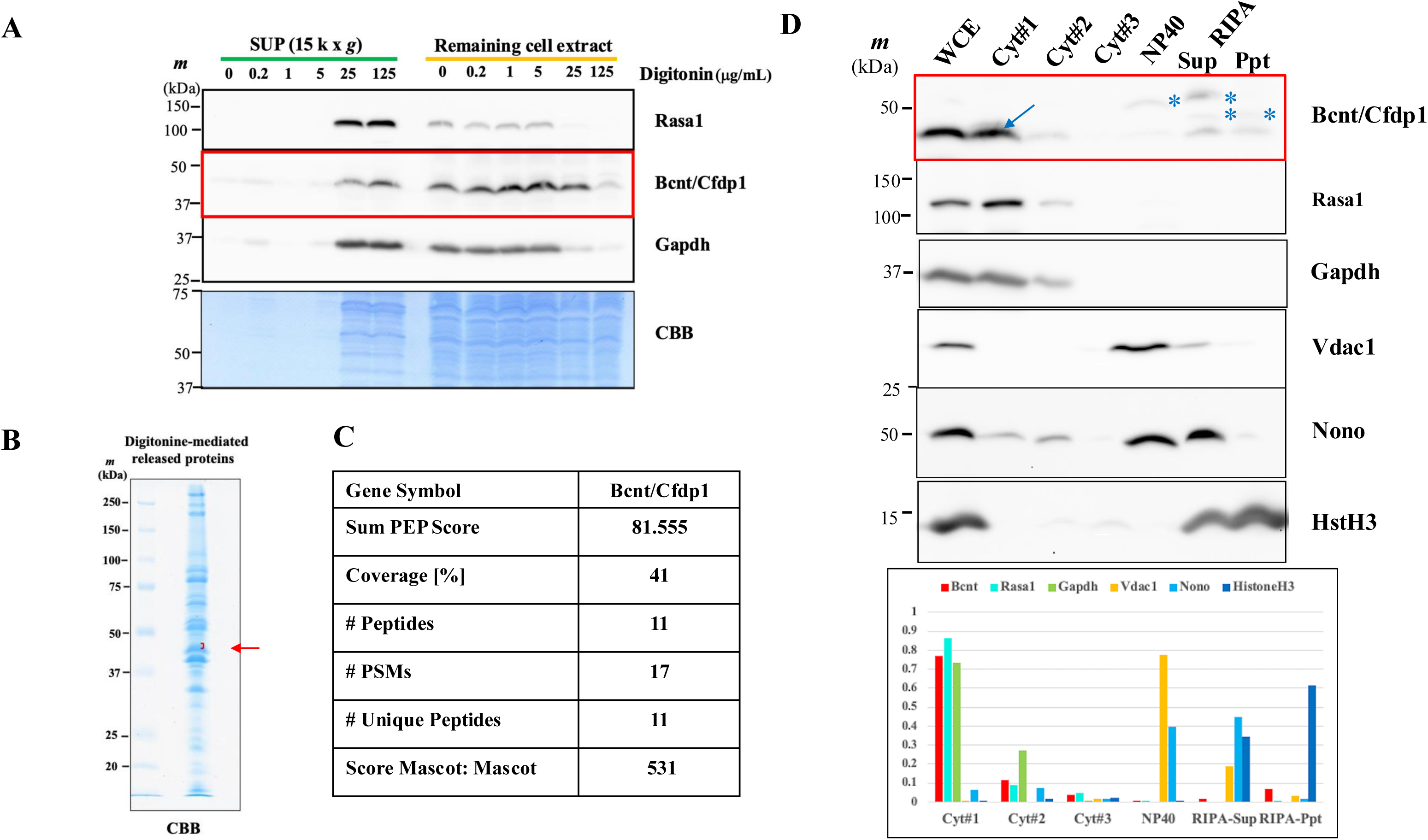
Release of Bcnt/Cfdp1 by digitonin treatment in an isotonic solution. **A.** Cell cultures in 12-well dishes were exposed to a range of digitonin concentrations in an isotonic solution for 10 min on a rocking shaker. After the supernatant of each eluate was collected, the corresponding extracts of the residual cellular debris were scraped in an SDS-containing buffer and boiled. Equal volumes of each sample were subjected to Western blotting using Abs against Rasa1 (upper box), Bcnt/Cfdp1 (middle box), and Gapdh (lower box). After Western blotting analysis, the membrane was stained with CBB to show the 37– 75 kDa region (at the bottom). **B.** The eluate supernatant was subjected to SDS/PAGE, and after staining the gel with CBB, the ∼43 kDa region was excised for mass spectroscopic analysis. **C.** After destaining the gel pieces, they were subjecting tryptic digests to liquid chromatography-tandem mass spectrometry analysis. Of the 124 proteins identified (Supplementary Table S4), data are shown for Bcnt/Cfdp1, with 41% recovered by the analyzed fragments. **D.** Using the three-step preparation of the digitonin-mediated eluate, as illustrated in Figure 7, Cyt #1–3 fractions were obtained using digitonin at 125 μg/mL in an isotonic solution. Following TCA precipitation, each pellet was dissolved in LS buffer (containing 1% SDS), and relatively equal volumes of each fraction were subjected to Western blotting analysis, as described in the legend of Figure 3. The dark blue arrow and asterisks correspond to the “upper band” of Bcnt/Cfdp1 and probable off-target band(s). Densitometric analysis of the Western blotting is shown at the bottom. The data represent the relative occupancy of each of the six marker proteins in each fraction (Cyt#1–RIPA–Ppt), with the total amounts of each marker protein set at 1.0.

Furthermore, we developed a three-step strategy to enhance the collection of cytoplasmic components, as shown in Figure 6. The supernatant released from the digitonin-treated cells was initially collected, centrifuged, and labeled Cyt#1. Subsequently, the cell debris on the dish was gently scraped off, and the supernatant was collected by centrifugation (Cyt#2). Next, the supernatant was obtained by washing the pellet with an isotonic solution (Cyt#3). Following the concentration of each fraction using trichloroacetic acid (TCA), the proteins were dissolved in sodium dodecyl sulfate (SDS)-containing buffer. Equal relative volumes of each fraction were subjected to Western blotting analysis, as shown in Figure 3. The elution pattern of Bcnt/Cfdp1 was similar to that of the cytoplasmic soluble marker proteins, such as Rasa1 and Gapdh (Figure 5D).

**Figure 6.**
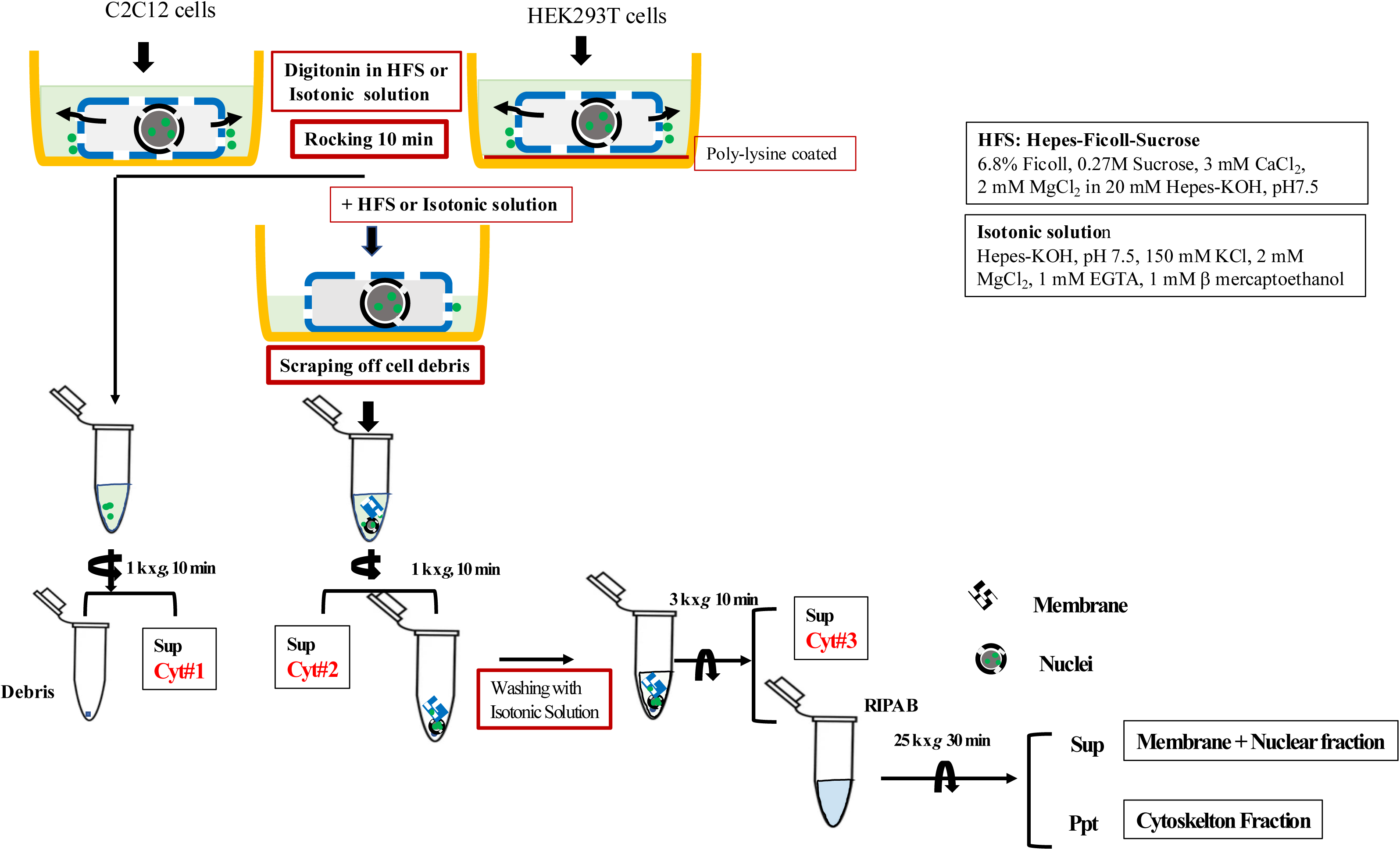
Schematic diagram of the three-step preparation of the digitonin-mediated eluates. Cultured cells were immersed in an isotonic or HEPES/Ficoll/sucrose (HFS) solution containing digitonin while the dish was rotated. The eluate supernatant was referred to as Cyt#1 after centrifugation. The remaining cell debris was scraped off using a cell scraper in a digitonin-free isotonic or HFS solution, and the supernatant was stored as Cyt#2 after centrifugation. The pellet was washed by suspension in an isotonic solution, and the supernatant was stored as Cyt#3 following centrifugation. The pellet was resuspended in RIPA buffer, sonicated, and centrifuged. The supernatant was used as the RIPA fraction, the pellet was dissolved in SDS-containing buffer, and the supernatant was analyzed as the Ppt fraction. In the case of isotonic solution, the pellets were solved in 0.2% Nonidet-40. The details of the procedure are described in the Materials and Methods section.

#### B) In Ficoll/sucrose solution

Macromolecular crowding has been shown to affect protein properties in living cells [2, 45], nuclear structure and function in preparation [46], and subcellular distribution in subcellular fractionation [47]. Therefore, digitonin treatment was performed in the HEPES-buffered Ficoll/sucrose solution (HFS) instead of an isotonic solution to prepare the cytoplasmic components from the cells appropriately. Cyt#1–3 fractions were prepared using the above three-step process (Figure 6) and subjected to Western blotting analysis. Notably, the elution pattern of Bcnt/Cfdp1 differed from that of the cytoplasmic soluble marker proteins (Figure 7A). A significant quantity of Bcnt/Cfdp1 was detected within the Cyt#3 fraction, whereas only a tiny amount was observed within the Cyt#1 and #2 fractions at lower digitonin concentrations. A comparative preparation was conducted using direct digitonin-mediated release in the HFS solution on HEK293T cells grown on polylysine-coated dishes (Figure 7B). The distribution pattern of BCNT/CFDP1 in HEK293T cells was similar to that observed in C2C12 cells. A significantly higher concentration of BCNT/CFDP1 was also observed in the Cyt#3 fraction than in the Cyt#1 and #2 fractions (Figure 7A). These results indicated that macromolecular crowding affects the mobility of Bcnt/Cfdp1 in C2C12 and HEK293T cells during the three-step preparation, suggesting that Bcnt/Cfdp1 exhibits flexible behavior in the process.

**Figure 7.**
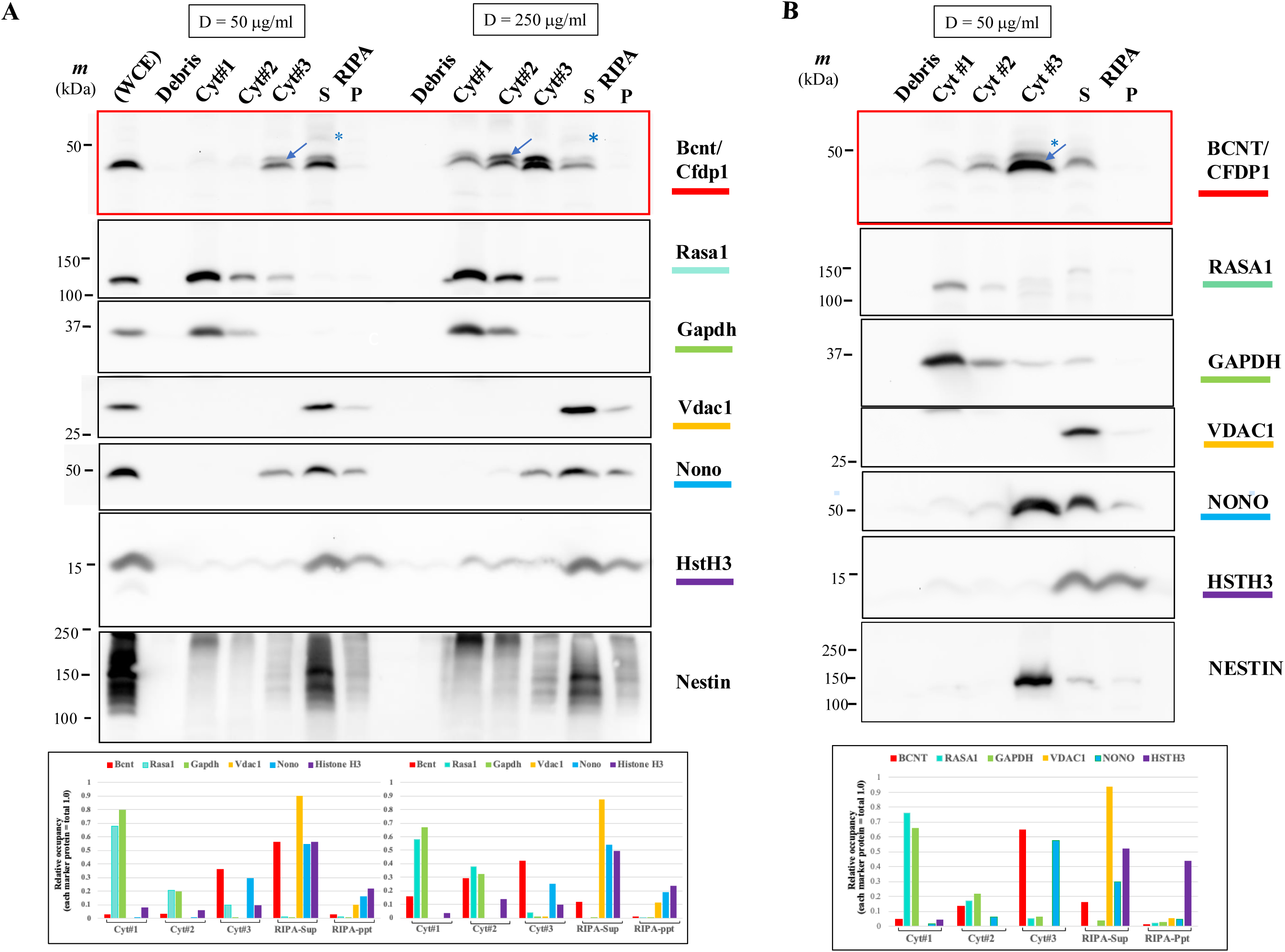
Differential release of Bcnt/Cfdp1 from cytosolic proteins by digitonin treatment in HFS. The Cyt#1–3 fractions were obtained using digitonin in HFS solution at 50 or 250 μg/mL in C2C12 cells or 50 μg/mL in HEK293T cells, as illustrated in Figure 6. Following TCA precipitation, each pellet was dissolved in LS buffer, and relatively equal volumes of each fraction were subjected to Western blotting analysis, as described in the legend in Figure 3. Densitometric analysis of the Western blotting is shown at the bottom. The data represent the relative occupancy of each of the six marker proteins in each fraction (Cyt#1–RIPA–Ppt), with the total amounts of each marker protein set at 1.0.

### Mobility of Bcnt/Cfdp1 in the Cytoplasm

We expected that the differential elution pattern of digitonin-mediated release in the HFS solution would reflect the mobility or integrity of each protein in the above three-step preparation. To examine the perspective of the property, we prepared Cyt#1 and #2* similarly to Figure 7, except that the pellet of Cyt#2 was washed with HFS solution instead of isotonic solution, and the washed supernatant was combined and designated as Cyt#2* (Figure 8A). These two fractions were subjected to quantitative LC-MS/MS analysis, and 2472 proteins were identified, of which 2442 were qualified for ratio evaluation (Supplementary Table S5). Among them, 51 glycolysis/glucogenesis- and TCA-related enzymes were listed, and their Cyt#1 to Cyt#2* ratios (Cyt#1/2*) were plotted as LOG2 (Figure 8 B and C, and Supplementary Table S6). The ratios of glycolysis-related proteins were above 1 (>0 in LOG2 unit), but TCA cycle-or membrane-associated proteins had ratios significantly below 1 (< 0 in LOG2), while the Bcnt/Cfdp1 ratio was 0.17 (-2.55 in LOG2) (Supplementary Table S5). Remarkably, the four isoenzymes of cytoplasmic and mitochondrial types revealed clear property trends.

**Figure 8.**
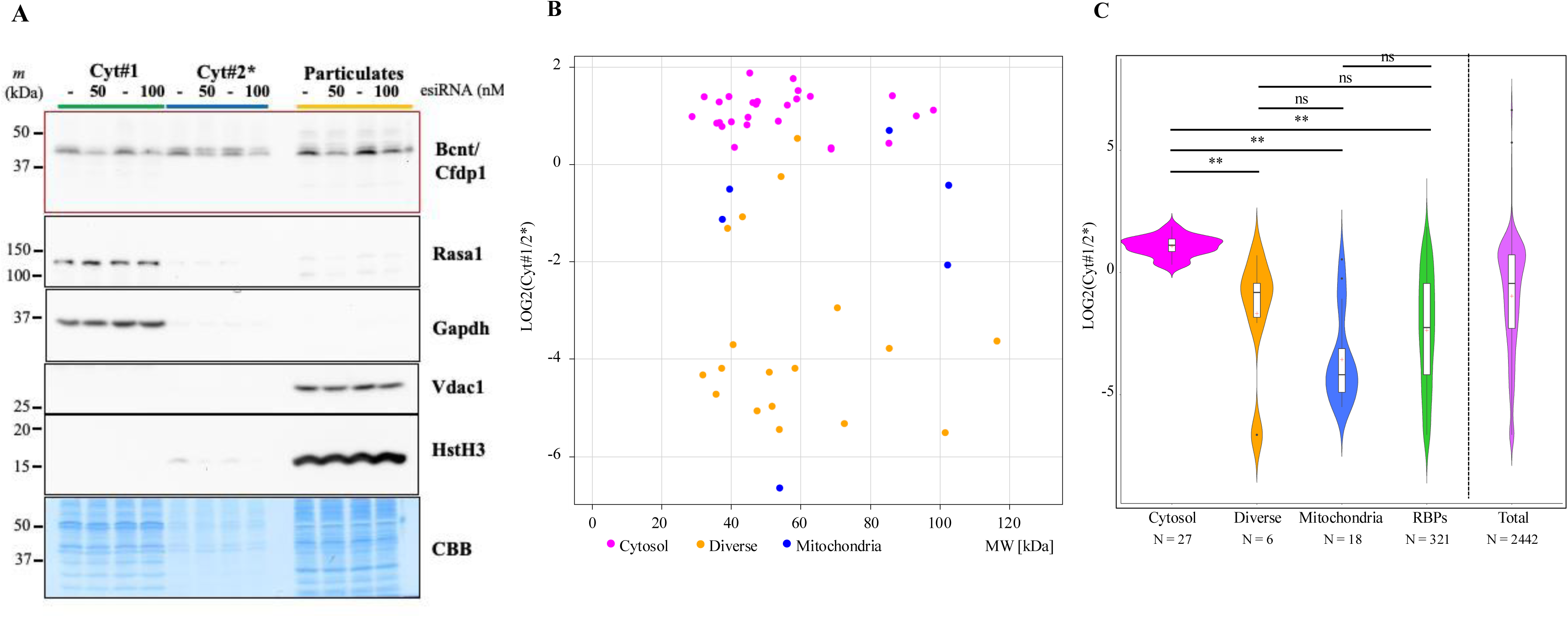
Mobility or integrity of digitonin-mediated released proteins in the Cytoplasm -Evaluation from the ratios of two fractions-. **A.** C2C12 cells treated with esiRNA (0, 50, or 100 nM) were subjected to digitonin-mediated fractionation in HFS solution at 250 µg/mL digitonin, as shown in Figure 6. But, after obtaining Cyt#2, its pellet was washed with HFS solution but not with isotonic solution, and the washed supernatant was combined to give Cyt#2*. Furthermore, the precipitate was dissolved in LS buffer after washing with HBS as a Particulate fraction. Equal amounts of protein from Cyt#1 (15 μg), Cyt#2* (7 mg), and Particulate (25 mg) fractions were subjected to Western blot analysis. These different protein amounts were adjusted so that the relative occupancy volumes of the three fractions were approximately equal. Each Cyt#1 and #2* fraction was subjected to quantitative mass spectrometry analysis (Supplementary Table S5). **B.** Fifty-one glycolysis/glucogenesis/TCA cycle/organelle-associated enzymes were listed from the 2442 qualified proteins by Buantitative mass analysis (Supplementary S6). Each Cyt#1/2* ratio value is plotted on the vertical axis as LOG2, while the horizontal axis shows its molecular weight (Mw). Glycolysis-related cytosolic proteins (magenta circles), organelle-related proteins (Diverse, orange circles), and TCA-related mitochondrial proteins (dark blue circles) are shown. **C.** The distribution of Cyt#1/2* ratios of 51-enzymes shown in Figure 8B (magenta, orange, dark blue), 2442 qualified total proteins (purple), and 321 RNA-binding proteins (green), which are selected based on published data were presented (Supplementary Table S7) as violin plots, respectively, using EZR software. The rectangles within the violin plots indicate the first and third quartiles, with central + as mean values. Results of statistical analysis are shown. ** indicates a significant difference with P<0.01

**Table 1.** Ct#1/2* ratios of four representative glycolysis/TCA-related isomers.

| Protein Names | Gene Symbol (Alias) | Intracellular localization | Mw (kDa) | Cyt#1/2* (Ratios) | Cyt#1/2*(LOG2) |
| --- | --- | --- | --- | --- | --- |
| Bucentaur/Craniofacial development protein 1 | Bcnt/Cfdp1 (cp27) |  | 33.4 | 0.17 | -2.56 |
| Aconitase | Aco1 | cytoplasmic | 99 | 2.27 | 1.18 |
|  | Aco2 | mitochondrial | 85.4 | 0.072 | -3.80 |
| Enolase | Eno1 | cytoplasmic | 47.1 | 2.36 | 1.24 |
|  | Eno2 | mitochondrial | 37.2 | 0.055 | -4.18 |
|  | Eno3 | cytoplasmic | 56.1 | 2.33 | 1.22 |
| Aspartate aminotransferase | Got1 | cytoplasmic | 46.2 | 2.41 | 1.27 |
|  | Got2 | mitochondrial | 47.4 | 0.03 | -5.06 |
| Malate dehydrogenase | Mdh1 | cytoplasmic | 40 | 1.835 | 0.88 |
|  | Mdh2 | mitochondrial | 35.6 | 0.038 | -4.72 |

Furthermore, the ratios Cyt#1/2* of 321 RNA-binding proteins selected based on published data were significantly smaller than that of the entire population (p-value <0.01) (Figure 8C and Supplementary Table S7). These results support the differential elution patterns of each protein and reflect the mobility or integrity of each protein during the three-step preparation, which is affected by macromolecular crowding.

## DISCUSSION

Although the lack of high-quality anti-Bcnt/Cfdp1 antibodies made the analysis difficult in this study, reliable data were obtained by quantitative mass spectrometry. Nuclear fractions and extracts of differentiated myotubes of C2C12 cells showed several off-target bands of Western blots using A305-624A-M as the anti-CFDP1 antibody [33], which is currently considered the most reliable antibody in our Western blot analysis. These difficulties forced us to focus on the cytoplasmic soluble proteins, which did not allow us to take advantage of the characteristics of C2C12 cells. In this study, cytochemical approaches showed that Bcnt/Cfdp1 was mainly observed in the nucleus, whereas biochemical fractionation detected the molecule at high levels in the cytoplasmic fraction. Here, we discuss these contradictory results from the viewpoints of nuclear export trafficking and macromolecular crowding and also address the potential functionality of Bcnt/Cfdp1 in the Srcap-C chromatin frame.

### Bcnt/Cfdp1 Subcellular Distribution Determined by Cytochemical Approaches

As reported in a previous study, we showed that tag-free mouse Bcnt/Cfdp1 is predominantly localized in the nuclei of human HEK293T cells using antibodies against mouse Bcnt/Cfdp1 peptides (Figure 1A, modified from Nakashima *et al*. [37]). This strategy was excellent, but these antibodies react with off-targets in Western blotting in HEK293T and C2C12 cells. Therefore, we used FLAG- and fluorescent protein-tagged Bcnt/Cfdp1 to assess its subcellular localization. The results showed that the signal was predominantly observed in the nuclei of C2C12 myoblasts and differentiated myotubes (Figure 1B-1 and 1C). Furthermore, using higher-resolution imaging in myoblast, Bcnt/Cfdp1 was detected in a segregated pattern, the regions with the lower DAPI staining (Figure 1B-2). The result is consistent with the observation that the *Drosophila* homolog YETI has been mapped to multiple sites on the euchromatic region on salivary gland polytene chromosomes [11].

When SRCAP-C is prepared at a higher-salt extraction buffer such as 300 mM KCl, BCNT/CFDP1 is scarcely captured as its component among other subunits [13, 48], which is, in part, why Bcnt/Cfdp1 has often been neglected [11]. However, a recent study revealed that BCNT/CFDP1 was significantly recovered when a lower-salt extraction buffer (150 mM NaCl) was used for isolation from the reconstituted system composed of recombinant proteins (personal communication, Dr. Shinya Watanabe, University of Massachusetts Medical School). This finding suggests that either weak ionic interactions are critical for BCNT/CFDP1 to be a component of the SRCAP-C or that high salt concentrations disrupt the specific environment, such as a condensate, in which Bcnt/Cfdp1 is incorporated into SRCAP-C. The N-terminal peptide of SWC5 was shown to have a potential for condensate formation through molecular dynamics simulations [49]. Although the amino acid sequence of the N-terminal regions is quite different between SWC5 and BCNT/CFDP1, both are IDRs rich in negatively and positively charged amino acid residues. Thus, we explored the possibility of condensate formation of BCNT/CFDP1 in the nucleus. The IDR of BCNT/CFD1 (the N-terminal 273 of 299 whole amino acids) was inserted into an optoDroplet assay vector containing a nuclear localization signal, and condensate formation was monitored [50] (Supplementary Figure S1). However, no positive results were obtained.

Considering the findings of the higher-resolution imaging in which Bcnt/Cfdp1 is detected in the lower chromatin density regions and the fact that the interchromatin is a porous medium [6], Bcnt/Cfdp1 may dynamically behave in the nucleus for coupling with Srcap-C. Further insights into the physical properties of Bcnt/Cfdp1 are expected in the context of weak ionic interactions in Srcap-C formation.

Nonetheless, considerable concerns remain with respect to the cytochemical results. First, we could not correctly evaluate the weak fluorescent signals in the cytoplasm, which were observed at the level of fluorescence microscopy. However, it was challenging to visualize after paraformaldehyde (PFA) fixation and subsequent anti-FLAG antibody detection, which are used to minimize artifact signals. Therefore, to accurately evaluate these weak signals, considering the side effects of exogenous protein expression, improvements in fixation and visualization methods, including expression systems that can control gene copy numbers, are necessary. Second, the fluorescent protein tag, corresponding to a mass of approximately 27 kDa, may inhibit passive nuclear export [8], resulting in overestimating Bcnt/Cfdp1 nuclear localization. Thus, it remains essential to prepare higher-quality anti-Bcnt/Cfdp1 antibodies.

### Effectiveness and Limitations of Western Blotting Monitoring

We employed A305-624A-M as an anti-Bcnt/Cfdp1 antibody to effectively monitor the ∼43 kDa band in Western blotting. The presence of Bcnt/Cfdp1 molecules in the ∼43 kDa region was confirmed by enrichment followed by mass spectrometry (Supplementary Figure S2 and Supplementary Table S1). However, in contrast to the cytoplasmic soluble proteins, the particulates and the extract of differentiated C2C12 myotubes showed several off-target bands in the wide range of ∼ 50 kDa (Figures 2C, 4, and 8). We reported similar bands in the precipitate fraction of T-Rex cell extracts solubilized in Radio-Immunoprecipitation Assay (RIPA) buffer (Supplementary Figure S4 in Iwashita *et al*. [33]). During myotube differentiation, another signal of ∼ 50 kDa gradually increased, whereas Bcnt/Cfdp1 levels decreased in the later stages (Figure 2E and Supplementary Figure S3). The latter was consistent with mRNA expression levels [51] (courtesy of Dr. Yan Fei Gao). Concerning the ∼43 kDa doublets, which are caused by phosphorylation in T-Rex cells (a HEK293 cell derivative) at the 250th serine residue of BCNT/CFDP1 [10] or the 245th serine residue of mouse Bcnt/Cfdp1 [33], its intensity in C2C12 cells was too weak to be accurately assessed. Taking these results together, we focused on their cytosolic extracts without mentioning the doublets.

### Subcellular Fractionation Using the Available Fractionation Kit

We first performed subcellular fractionation using a Thermo Scientific kit containing multiple non-ionic detergents in each step [38]. Using the process, significant bands of 43–45 kDa were observed in the cytoplasmic fractions of both C2C12 and HEK293T cells, and other bands of ∼50 kDa were detected in the respective nuclear fractions (Figure 3). The presence of Bcnt/Cfdp1 in the cytoplasmic fraction was confirmed via mass spectrometry (Figure 5C from Supplementary Table S2). Non-ionic detergents are often combined with a hypotonic solution to achieve optimal results in sequential subcellular fractionation and have been reported to be particularly effective in separating the cytoplasmic and nuclear compartments [52]. However, the possibility that non-ionic detergents, even at low concentrations, affect lipid-lipid and lipid-protein interactions, affecting the membrane-associated population of the targets and compromising membrane integrity cannot be ignored.

The nucleus is protected by a fibrous layer called the lamina, which lies beneath the nuclear envelope [53]. Therefore, the apparent intactness of the nucleus after various treatments does not necessarily guarantee the integrity of the nuclear envelope. The nuclear membrane is much more elastic than previously thought. Its integrity, even its rupture, is influenced by a physiological response [54] and through signaling pathways, such as the activation of Akt kinase [55]. Regarding the distribution of Bcnt/Cfdp1, it is interesting that HMGB1 was detected in the cytoplasm using the non-ionic detergent IGEPAL (Nonidet P-40), but in the nucleus using the Dounce homogenization method in lung cancer cell lines [56]. The discrepancy may be attributed to the properties of HMGB1 that include nuclear localization signals in the N-terminal and IDRs in the C-terminal. Notably, blocking the nuclear localization signals of HMGB1 in leucocytes redirects it to the cytoplasm, eventually leading to secretion [57]. In our mass analysis, Hmgb1 was mainly detected in the cytoplasmic fraction in the subcellular fractionation of C2C12 and annotated in most compartments, including the non-structural extracellular matrix (Supplementary Table S2). HMGB1, found initially as a nuclear protein, may have properties similar to those of Bcnt/Cfdp1 in flexible intracellular behavior. Regarding the possibility of Bcnt/Cfdp1 secretion, it was reported as an unconventionally secreted protein in the conditioned medium of C2C12 myotubes [58]. Furthermore, a secretome prediction tool supported the result: OutCyte, http://www.outcyte.com/(courtesy of Dr. Gereon Poschmann) [59]. Although we tried to confirm the possibility, obtaining the C2C12 cell culture medium supernatant was difficult even in myoblast cultures, as dead cells constituted at least 1% of the total. Thus, it was impossible to reach a definitive conclusion, and further research is needed.

### Digitonin-Mediated Extraction in Isotonic or Ficoll/Sucrose Solution

To examine Bcnt/Cfdp1 in the cytoplasm more strictly, we directly prepared the cytoplasmic soluble proteins from cell cultures on the dishes by digitonin treatment. The method was expected to allow the extraction of cytosolic proteins with minimal interference from other proteins. In the experiment, Bcnt/Cfdp1 was detected along with the cytoplasmic soluble proteins Rasa1 and Gapdh in isotonic buffer at a concentration of 25 μg/mL (0.0025%) (Figure 5A and Supplementary Table S4). Notably, a nuclear protein, Trip13 (thyroid hormone receptor interactor 13), was released with the digitonin treatment and showed similar mobility to Bcnt/Cfdp1 (Supplementary Table S2 and S4). Indeed, the human ortholog TRIP13 has been reported to be significant trafficking in the cytoplasm and the nucleus [44]. These results support the cytoplasmic distribution of Bcnt/Cfdp1 and the specific proteins that exhibit similar molecular mobility. On the other hand, it was difficult to obtain reproducible release patterns at lower digitonin concentrations because of the tailing of cytoplasmic soluble marker proteins under the direct elution method, especially in the HFS solution.

Since macromolecular crowding is critical for molecular behavior [2, 45], we referred to the following reports to examine the subcellular behavior and distribution of Bcnt/Cfdp1. First, solutions without macromolecular crowding may not fully preserve the nuclear structure and function during nuclear isolation [46], and second, the intracellular trafficking of SIRT1 may depend on macromolecular crowding [47]. Therefore, according to the report, we used the HFS solution instead of an isotonic solution in the digitonin treatment [47].

Notably, Bcnt/Cfdp1 was distinctly extracted compared to the cytoplasmic soluble proteins, especially Rasa1, appearing in the Cyt#3 fraction rather than in the Cyt #1 and Cyt #2 fractions (Figure 7). Cyt#3 was the supernatant fraction obtained by scraping the digitonin-treated cell debris and washing the pellet with an isotonic solution, followed by centrifugation (Figure 6). A comparable extraction pattern was also observed at a higher digitonin concentration of 250 μg/mL in C2C12 cells and 50 μg/mL in HEK293T cells.

We prepared the fractions of Cyt#1 and Cyt#2*, the latter obtained by washing the pellet of Cyt#2 with HFS solution and combining the washed supernatant (Figure 8). The mass analysis of Cyt#1 to #2* of ratios reflected the mobility of each protein in the cytoplasm. The ratio of Bcnt/Cfdp1 was 0.17 (-2.55 in LOG2), suggesting that Bcnt/Cfdp1 exists in a complex in the cytoplasm, probably associated with RNA or RNA-binding proteins [28]. This result reflects an essentially similar phenomenon, as reported; the subcellular distribution of SIRT1 differs depending on whether the fractionation solution is composed of macromolecular densities [47]. Although the rate at which the nuclear membrane is partially damaged, resulting in leakage out of the nucleus during the three-step preparation, should not be underestimated, we propose that Bcnt/Cfdp1 and some specific proteins have physical properties that allow them to be readily mobile from the nucleus to the cytoplasm, which is affected by macromolecular crowding. These findings have practical implications for future research and applications in protein behavior.

### Beyond the Limitations of Digitonin Treatment **-**Conflicting Results for the Subcellular Localization of Bcnt/Cfdp1

The discrepancy in the subcellular distribution of Bcnt/Cfdp1 described above is a complex and vital issue that presents significant challenges. As mentioned earlier, several factors could lead to discrepancy. Two situations lead to overestimating nuclear localization: the cytochemical approach could not capture the weak signals in the cytoplasm, and the fluorescent protein tag could inhibit passive nuclear export. On the other hand, mobility due to nuclear membrane damage in the digitonin-mediated preparation could lead to the overestimation of cytoplasmic localization.

It was challenging to completely recover the cytoplasmic soluble proteins released by direct digitonin treatment, especially in an HFS solution. Subsequently, we developed a three-step strategy for the preparation of digitonin-mediated extracts. The Cyt#3 fraction obtained by washing the digitonin-mediated cell debris pellet contains the proteins of the residual cytoplasmic population, which may include the attached one to various organelle membranes. In addition, proteins present in the nucleus may leak out of the nucleus due to damage to the nuclear membrane during the three-step procedure. However, given the relatively selective activity of digitonin in membrane poration, which depends on the cell type and physiological state, it is inherently challenging to avoid partial membrane damage during digitonin treatment to harvest the cytoplasmic components completely. For example, although digitonin treatment can be mild, i.e., using 0.001% in phosphate-buffered saline (PBS) for 2 min on ice, it still caused partial damage to the nuclear envelope, as evidenced by staining with an anti-lamin B1 Ab [60]. In addition, digitonin is capable of disordering the alkyl chains of phospholipid monolayers, and its effect is more potent on phosphatidylethanolamine and phosphatidylserine than phosphatidylcholine [61]. Orre *et al*. performed a large-scale analysis of subcellular localization in five cancer cell lines using a uniform fractionation method that involved the digitonin treatment of cell cultures on dishes, followed by homogenization in a hypotonic buffer [41]. Whereas the data showed 80% agreement between the cell lines, significant discrepancies were attributed to the differences in the localization of proteins, including BCNT/CFDP1, between the cytosol and nucleus (https://lehtio-lab.se/subcellbarcode/). This discrepancy might be due to the different susceptibility of the nuclear envelope in some cells than in others during the fractionation or due to other factors, such as differences in the levels of post-translational phosphorylation in each cell. Furthermore, the unique intracellular mobility that we observed on Bcnt/Cfdp1 and some proteins may also contribute similarly.

Beyond these difficulties of evaluation in digitonin-mediated preparation, our data suggest that Bcnt/Cfdp1 and some proteins are readily mobile from the nucleus. Recent studies have revealed unique features for the nuclear membrane compared to other cell membranes; nuclear lipid membranes are much more elastic than other membranes [62], and their lipid composition is closely related to membrane integrity [63]. Additionally, NPC, the sole transport gateway between the cytoplasm and nucleus, comprises approximately 300 IDPs, rendering the pore a flexible rather than a rigid structure. Interestingly, while folded protein transport has revealed a size-dependent exclusion, where larger proteins are restricted, IDPs can diffuse through the NPC without energy or specific transport receptors due to their lack of a fixed structure, allowing them to navigate the pores more efficiently [9]. These results support the possibility that Bcnt/Cfdp1 and some specific proteins may be able to diffuse rapidly through the NPC into the cytoplasm during the subcellular fractionation. However, leakage due to partial damage to the nuclear envelope cannot be underestimated in the three-step processes.

### Biological Functionality of Bcnt/Cfdp1

*Bcnt/Cfdp1* plays a pivotal role in developmental processes, as evidenced by its mutants in *Drosophila* [20], zebrafish [21, 22], and mice [23, https://www.jax.org/strain/033834]. These biological variations include alterations to the cell cycle, cell migration, and apoptosis, which subsequently influence cancer prognosis and susceptibility [28, 29].

The zebrafish mutant *gazami*, lacking the seventh exon of Bcnt/Cfdp1 (encoding the C-terminal 31 amino acid residues), exhibited abnormal neuronal differentiation in brain granule cells. The elevated expression levels of cyclin B1 (*Ccnb1*) have been shown to contribute to inappropriate differentiation [21]. However, its upregulation might be one of the gene expression variations induced by deficient *cfdp1*, similar to the *Bcnt/Cfdp1* knocked down in the Cfdp1-K1 cells [33]. Interestingly, many genes related to the Wnt or Akt pathways and apoptosis/autophagy, such as *Wls*, *Wnt4*, *Anxa2*, *Bcl2*, and *Akt2,* are included, up- or downregulated in Cfdp1-K1 cells (Supplementary Table S8). They are involved in cell cycle control, cell migration, and apoptosis. Indeed, the Wnt signaling pathway is suppressed specifically in extraluminal mesenchymal cells of the valves of zebrafish Bcnt/Cfdp1-deficient embryos [22].

The broad effects of deficient *Bcnt/Cfdp1* may be comparable to natural genetic variations in the fission yeast *swc5*. The mutant exhibited abnormal mRNA, antisense, and noncoding RNA expression, possibly caused by improper histone exchange reactions [34]. Srcap-C has been demonstrated to be involved in differentiation processes in intestinal epithelial cells controlled by H2A.Z in association with Wnt signaling [64]. Assuming that Bcnt/Cfdp1, similarly to SWC5 in SWR1-C, is a key factor for Srcap-C activity [14–16], a deficiency in ***Bcnt/Cfdp1*** may induce inappropriate histone exchange reaction activity, resulting in changes to the transcriptional landscape by Srcap-C. The phenotype of the mutant *gazami* described above may be a clear example.

Notably, some reports of the biological responses of perturbed *BCNT/CFDP1* mentioned earlier are partially associated with *BCAR1,* a proximal neighboring gene of *BCNT/CFDP1* located on chromosome 16q23.1 and has multiple functions [65]. The *BCAR1–CFDP1–TMEM170A* locus, which is highly conserved across vertebrates, contains several single nucleotide polymorphisms (SNPs) associated with diseases, such as vascular diseases [66] and Dupuytren’s disease [67]. Although these diseases may be highly related to variations in the Wnt/Notch/Akt signaling pathways, the causal relationship between these SNPs and diseases remains unclear.

## CONCLUSION

Metazoan Bcnt/Cfdp1 is predicted to have a similar function to yeast SWC5 as a component of Srcap-C. Consistent with this prediction, both untagged mouse Bcnt/Cfdp1 transfected into HEK293T cells and fluorescent protein-tagged mouse Bcnt/Cfdp1 in C2C12 cells were detected primarily in the nucleus by cytochemical approaches. However, by detergent-based biochemical fractionation of C2C12 and HEK293T cells, endogenous Bcnt/Cfdp1 was detected mainly in the cytoplasmic fraction. Furthermore, when cell cultures on the dish were treated with digitonin, Bcnt/Cfdp1 was released and detected as a cytoplasmic protein, while different elution patterns were observed between isotonic and Ficoll/sucrose solutions. Although our results on the intracellular behavior of Bcnt/Cfdp1 reveal its mobility in the presence of detergent rather than the export trafficking, we argue that Bcnt/Cfdp1 and some specific proteins have physical properties that allow them to be readily mobile from the nucleus to the cytoplasm, which is affected by macromolecular crowding. The unique property of Bcnt/Cfdp1 intracellular mobility remains unclear, but its elastic properties at the cytoplasm/nucleoplasm interface may be relevant. These issues are the most critical aspects of this study and are expected to become more evident as our understanding of the nuclear membrane expands beyond classical cell membrane studies [68].

### Data Availability Statement

All data reported in this paper, including the proteomics mass spec dataset, are available without restriction.

## MATERIALS AND METHODS

Supplementary Table S9. provides detailed information on the material resources used in the study, except for some experiments mentioned therein.

### Cell Culture

C2C12 cells were obtained from the European Collection of Authenticated Cell Cultures through KAC Co., Ltd. (Kyoto), expanded, and aliquots were stored at –80°C using Bambanker. HEK293T cells were provided by Dr. Rie Komori (Tokushima Bunri University) and stored in liquid nitrogen using Cellbanker. Both types of cells were cultured in Dulbecco’s Modified Eagle’s Medium (DMEM)–10% FCS and subcultured using Accutase, as described previously [33]. Briefly, after incubation for 5 min at room temperature, the cells were gently homogenized using a 1000 μL tip, mixed with medium, centrifuged, and resuspended in the medium unless otherwise specified. The cell numbers were counted using a hemocytometer (a disposable cell counting plate of the Improved Neubauer Type), and dead cells were estimated via the trypan blue exclusion assay. Before seeding, the dishes were incubated with the medium in a CO_2_ incubator for at least 30 min.

### C2C12 Differentiation

C2C12 cells (5 x 10^4^ per 35 mm dish) were cultured for 2 days, and then the medium was replaced with DMEM–2% horse serum (designated as day 0). On the corresponding day, the cultures were washed with chilled HEPES buffered saline (HBS) twice and homogenized in 100 μL Lysis buffer (LS:1% SDS, 1 mM EDTA in 10 mM HEPES–NaOH, pH 7.5) supplemented with inhibitors of proteases and phosphatases using a silicon policeman. Each extract was collected in a 1.5 ml tube, boiled immediately for 3 min, and then sonicated for 2.5 min. A constant volume of each extract was subjected to 12.5% SDS-polyacrylamide gel electrophoresis (SDS/PAGE), followed by Western blot analysis.

### Coating Culture Dishes with Polylysine

According to the protocol, 60 mm dishes were coated with polylysine with some modifications (https://assets.thermofisher.com/TFS-Assets/BID/Flyers/gibco-poly-d-lysine-flyer.pdf). Each dish was shaken with 1 mL of 50 μg/mL poly-d-lysine solution on a seesaw shaker for 1 h at room temperature. To examine fluorescent protein-tagged mBcnt/Cfdp1, a commercially available 4-well polylysine-coated chamber (Matsunami, Osaka) was used.

### Construction of Plasmids

All Polymerase Chain Reaction (PCR) primers are listed in Supplementary Table S10. Tag-free and FLAG-tagged Bcnt/Cfdp1 were generated as previously described [37]. The same primers were used to generate mCherry- and EGFP-tagged MCS or mouse Bcnt/Cfdp1 using their respective expression vectors (NCBI accession numbers LC311025, LC311020, LC311024, and LC311019).

### Transfection of Plasmids

Transfections were carried out using Lipofectamine 3000 according to the manufacturer’s protocol with minor modifications. Prior to transfection, each half of the medium was saved once, and the plasmids were transfected. After 4 h, the saved medium was returned, and the cells were cultured for 20-44 h. For the fluorescent protein observation, C2C12 cells were seeded at a density of 1 × 10^4^ cells per well in 0.75 mL medium per well of polylysine-coated chamber, which was pre-incubated with culture medium in a CO_2_ incubator. After 20 h culturing, the medium was replaced with 0.5 mL of antibiotic-free medium, and cells were transfected with 0.25 μg of each plasmid using Lipofectamine 3000 at a DNA-to-Lipofectamine ratio of 1:2. Following 4 h of culture, 0.5 mL of medium containing 20% FCS was added to each well. The cells were then cultured for an additional 20 h.

### Fluorescent Proteins by Immunocytochemistry

The transfected and further cultured cells were washed twice with cold HBS and fixed with a final concentration of 4% PFA by dilution in 0.1 M phosphate buffer (pH 7.2). The cells were washed with 10 mM PBS to remove PFA. All the blocking serum, antibodies, and nuclear staining dye were used at the indicated dilutions with 10 mM PBS containing 0.2% Triton X-100. The cells were blocked with 5% normal donkey serum for 30 min at room temperature, then incubated with the anti-DYKDDDDK tag monoclonal Ab (1:1000) at 4°C overnight. The immunoreactivity was visualized with Cy3-conjugated donkey anti-mouse IgG (1:200) or CF™488A-conjugated donkey anti-mouse IgG (1:200). DAPI (1 μg/mL) was used for nuclear staining. The immunostained cells were examined by phase contrast and fluorescence microscopy (TE2000-U, Nikon).

### Enrichment of ∼43 kDa Band by Western Blot Monitoring

A puromycin-resistant cell line of AB2.2 mouse embryonic stem cells was gifted from Dr. Tsutomu Kumazaki (Tokushima Bunri University), who obtained the original AB2.2 cells from the Sanger Institute (Cambridge, UK) and generated the cell line by transfection with pMSCVpuro plasmid using PolyFect. The cells were expanded under serum- and feeder-layer-free culture conditions as described in [33] without G418 and puromycin. The cells were harvested using Accutase and rinsed with chilled HBS. The cell suspension aliquots were snap-frozen in liquid nitrogen and stored at −80°C until use. For the experiment, the cell suspension (equivalent to 3 x 10^7^ cells) was thawed and resuspended in 300 μL of Solution 1 (a cytoplasmic extraction buffer from Atto, Tokyo). The suspension was homogenized in a Biomasher II, rotating the pestle 10 times, sonicated in an ice-water bath (10 s, three times), and then centrifuged at 25,000 × *g* for 30 min at 4°C. Subsequently, CHAPSO (C_32_H_58_N_2_O_8_S) was added to the supernatant at a final concentration of 1% (w/v). The extracts were slowly loaded onto heparin-agarose (200 μL bed volume in a spin column) by cycling three times after the resin had been equilibrated with Solution 1. Nonidet-P40 was included in all buffers at a final concentration of 0.05% v/v, which was utilized for the subsequent chromatographic procedures. After washing the resin with 20 mM Tris-NaOH, pH 7.5, the bound proteins were eluted twice with 200 μL of Tris buffer containing different concentrations of NaCl. Each elution was allowed 5 min to settle at each elution step. The eluates from the heparin column chromatography with 0.05–0.1 M NaCl were combined and loaded onto Phos-tag resin (150-μL volume in a spin column) by three cycles. The resin was washed thrice with 150 μL of wash buffer, followed by a buffer containing 1M NaCl. The bound proteins were released using 4 × 100 μL of phosphate-containing elution buffer without Nonidet-P40 following the manufacturer’s protocol (https://labchem-wako.fujifilm.com/jp/category/docs/01072_doc01.pdf). The eluates were concentrated by deoxycholate-TCA precipitation, as described in a standard protocol (https://www.sigmaaldrich.cn/deepweb/assets/sigmaaldrich/product/documents/132/021/protprb <u>ul.pdf</u>). One aliquot of preparation was stored in a buffer designed for two-dimensional gel electrophoresis (Solution 2 from Atto Corp.). Following SDS/PAGE separation, the gel was stained in silver staining, and a gel piece in the range of ∼43 kDa was excised for mass spectroscopy analysis.

### Subcellular Fractionation using Available Fractionation Kit

C2C12 or HEK293T cells were harvested using Accutase, washed with DMEM-1% polyvinyl alcohol (PVA), and then washed with chilled PBS. Cells were fractionated according to the manufacturer’s protocol, with modifications to minimize the contamination for the next step (Thermo Fisher Scientific, https://assets.thermofisher.com/TFS-Assets/LSG/manuals/MAN0011667_Subcellular_Protein_Fraction_CulturedCells_UG.pdf). The samples were centrifuged, and the resulting supernatant was combined with the original solution. The nuclear extraction buffer without CaCl_2_ and nuclease was used for washing for chromatin-bound nuclear fractionation. Each fraction was concentrated by TCA precipitation, as described below. The resulting pellets were then dissolved in LS buffer and added to concentrated SDS/PAGE sample buffer at a final concentration of 1 x boiled for 3 min. Finally, equal relative volumes of each fraction were subjected to Western blotting analysis. To evaluate the resulting five fractions, Rasa1 and Gapdh were used as marker proteins for the cytoplasmic fraction, Vdac1 for the membrane fraction, NonO for the soluble nuclear fraction, and Histone H3 for the chromatin-bound fraction. For quantitative mass spectrometry analysis, C2C12 cells were seeded at 1.7 x 10^6^ per 100 mm dish x 4 and cultured for 23 h. Duplicate samples were prepared independently in subsequent steps. The cells were rinsed twice with 10 mL of warm HBS and treated with 2 mL of Accutase per 100 mm dish. The incubation was carried out at room temperature for 10 min with gentle rocking on a seesaw shaker while rotating the cell dish at 90 degrees per 2 min. The suspended cells were gently homogenized with a 1000 μL tip to prepare a single-cell suspension, which was then collected in a 15 mL tube. The cell suspensions were combined by washing the dishes twice with 2 mL of chilled DMEM-1% PVA and centrifuged for 1 min at 4°C. The percentages of trypan-blue-positive cells in the two groups were 1.8 and 2.2%, respectively. The cells were subjected to a subcellular fractionation at 10^6^ cells per 100 μL of cytoplasmic solution from the kit. Each subcellular fractionated sample was concentrated with TCA, as described above.

### Preparation of Digitonin Solution

A stock solution of digitonin was prepared by mixing 25 mg per mL of H_2_O at ∼80°C. The solution was then aliquoted and stored at –30°C. Before usage, the thawed aliquot was briefly sonicated, heated to ∼60°C, cooled to room temperature, and diluted as needed.

### Digitonin-Mediated Release in Isotonic Solution from C2C12 Cells

C2C12 cells (4 x 10^5^ cells) were seeded in a 6-well dish and cultured in 2 mL DMEM–10% FCS for 26 h. After two rinses with 2 mL of chilled HBS, the plates were transferred to a cold room, and sample preparation was performed below 4°C. The residual buffer was wholly removed with filter paper, and the cells were immersed in 0.5 mL of isotonic solution (HEPES– KOH, pH 7.5, 150 mM KCl, 2 mM MgCl_2_, 1 mM EGTA, 1 mM β-mercaptoethanol). The plate was gently rocking on a seesaw shaker for 10 min. Each fluid was collected into a 1.5 mL tube and centrifuged at 10,000 × *g* for 10 min at 4 °C. Concurrently, using a silicone policeman, each corresponding remaining cell in a 6-well dish was lysed in 100 μL of LS buffer, which included a cocktail of inhibitors against proteases and phosphatases. Extracts were collected into 1.5 mL tubes using a 200 μL bore-type tip. After vortexing and boiling for 5 min, the samples were sonicated for 2.5 min. Each sample was concentrated with TCA and dissolved in LS buffer using a mixer for over 1 h (Tomy Digital Biology, Tokyo). Final samples were prepared by mixing 2× SDS/PAGE sample buffer to adjust to 1 x.

### Subcellular Fractionation with Digitonin in a Three-Step Process

C2C12 cells (3 x 10^5^) or HEK293T cells (4 x 10^5^ cells in the polylysine-coated dish) were cultured in a 60 mm dish for 42–44 h. Washing was performed in the same way as described earlier. The cells were immersed in 1 mL of digitonin-containing isotonic solution or HFS solution (6.8% Ficoll, 0.27 M sucrose, 3 mM CaCl_2_, 2 mM MgCl_2_ in 20 mM HEPES–KOH, pH7.5) and gently rocked for 10 min while rotating the cell dish at 90 degrees per min on a seesaw shaker to ensure even distribution of the solution. The dishes were placed on ice, and the supernatant fluid was collected in a 1.5 mL tube. The tube was centrifuged at 1000 × ***g,*** 10 min, 4°C, and the resulting supernatant was designated Cyt#1, while the pellets were examined as debris. The remaining cell layer was gently scraped off into a 0.5 mL isotonic or HFS solution without digitonin using a cell scraper, and the extract was collected into a 1.5 mL tube using a bore-type 1000 μL tip. The sample was mixed by vortexing for 5 s at dial 8, left to settle for 15 min on ice, and then centrifuged at 1000 × *g* for 10 min at 4°C; the supernatant was saved as Cyt#2. The pellet was then homogenized in 500 μL isotonic solution. The homogenates were mixed by vortexing and gentle pipetting with a bore-type 1000 μL tip and left for 5 min on ice before centrifugation at 3000 × ***g*** for 10 min at 4°C. The supernatant was saved as Cyt#3. The pellets were resuspended in 150 μL of RIPA buffer (20 mM Tris-HCl, pH 7.5, 150 mM NaCl, 0.5% sodium deoxycholate, 1% Nonidet-P40, 1 mM EDTA) containing proteinase and phosphatase inhibitors. In some experiments, the pellets were treated with 0.2% Nonidet-P40 and analyzed as an NP40 fraction before being dissolved in RIPA buffer. Subsequently, they were sonicated in an ice-water bath for 2.5 min (10 s intervals, 15 times) and centrifuged at 25,000 × *g* for 30 min at 4°C. The supernatant was designated RIPA-Sup, while the pellets were dissolved in 50 μL LS. They were vortexed, immediately boiled for 3 min, and analyzed as the RIPA–Ppt fraction.

### Treatment with esiRNA

C2C12 cells were seeded in a 24-well plate at 2 x 10^4^ cells per well. After 20 h culture, the medium was replaced with 300 μL of either regular medium (A group) or FCS-free DMEM (B group). Four hours later, transfection was performed with 0, 50, or 100 nM of esiRNA targeting mouse *Cfdp1* (Sigma-Aldrich, dissolved in 1 mM sodium citrate, pH 6.4) using Lipofectamine 3000. After 4 h, 300 μL of regular medium (to A group) or DMEM containing 20% FCS (to B group) was added, and the cells were cultured for an additional 44 h. The cells were rinsed twice with chilled HBS and scraped off using a silicone policeman in 50 μL of LS buffer supplemented with inhibitors against proteases and phosphatases. The cell lysates were harvested in a 1.5 mL tube and then boiled immediately. After sonication (10-s intervals x 10), the samples were centrifuged at 25,000 × *g* for 30 min at 22°C. Constant amounts of protein (20 μg) were subjected to Western blot analysis. For examination of siRNA effects on subcellular fractionation, C2C12 cells were seeded at a density of 5 x 10^4^ or 1 x 10^5^/35 mm dish and cultured for 19 h, treated with esiRNA (0, 50, or 100 nM) using Lipofectamine for 44 h. The digitonin-mediated fractionation was carried out in an HFS solution similar to the above three-step process with the following modification. After obtaining the supernatant of Cyt#1 using 0.4 mL of an HFS solution containing 250 μg/ml digitonin, the residual cell layers were scraped off in 0.2 mL of an HFS solution and centrifuged without vortexing. The pellets were suspended in 0.2 mL HFS solution with brief vortexing and standby for 15 min on ice, then centrifuged, and the supernatant was combined as Cyt#2*. After washing the second pellets with HBS, the pellets were resolved in LS buffer, and the fraction was designated as Particulate. Cyt#1 and #2* were precipitated by TCA treatment, and the precipitates were resolved in 80 μL LS buffer. Equal amounts of proteins in each fraction of Cyt#1 (15 μg), Cyt#2*(7 μg), and Particulate (25 μg) were subjected to Western blot analysis.

### Examination of Potential LLPS of BCNT/CFDP1 in the Nuclei

Plasmid construction and optoDroplet assays were performed in the same way as previously reported [50]. The IDR of BCNT/CFDP1 was estimated using the PONDR VSL2 algorithm (http://www.pondr.com/). The IDR fragment of BCNT/CFDP1 (residues 1–273) was amplified by PCR from a human cDNA library (generated from total RNA extracted from HEK293T cells. Primers were described in Supplementary Table S10). An optoDroplet assay vector was constructed from the Cry2 vector and mCherry using the NEBuilder HiFi DNA Assembly Kit to incorporate the BCNT/CFDP1 fragment. HEK293T cells were transfected with the plasmids using TurboFect, and potential LLPS activity was measured after 48 h by image monitoring (Nikon Ti2 inverted microscope with the A1R confocal laser scanning system). After acquiring background mCherry images, cells were imaged using blue light stimulation (488 nm stimulation/3 s, interval/17 s, mCherry imaging/10 s x 3), followed by mCherry imaging (imaging/10 s and interval/50 s x 10).

### Isolation of Transiently Expressed FLAG-tagged Bcnt/Cfdp1

C2C12 cells (5 x 10^5^ cells/100 mm dish) were cultured for 21 h, transfected with the F-*Bcnt/Cfdp1* plasmid (10 μg), and cultured for an additional 42 h until they were nearly confluent. The cells were rinsed twice with chilled HBS on ice and scraped off using a cell scraper in 0.5 ml RIPA buffer containing inhibitors against proteinases and phosphatases. The lysates were collected in a 1.5 mL tube, sonicated for 2.5 min (15 × 10-s pulses at 10-s intervals), and centrifuged at 25,000 × *g* for 30 min at 4°C. The resulting supernatant was snap-frozen in liquid nitrogen and stored at −80°C until use. The frozen extract was thawed, sonicated in an ice-water bath (3 × 10-s pulses at 10-s intervals), and centrifuged at 10,000 × *g* for 1 min at 4°C. The isolation of F-Bcnt/Cfdp1 was essentially the same as described previously [33] except for using RIPA buffer to equilibrate the agarose before mixing the cell extract.

### TCA Precipitation

TCA precipitation was carried out according to a protocol with some modifications (https://www.thermofisher.com/document-connect/document-connect.html?url= https://assets.thermofisher.com/TFS-Assets%2FLSG%2Fmanuals%2FMAN0011667_Subcellular_Protein_Fraction_CulturedCells_UG.pdf). The pellets were washed once with acetone and twice with a 4:1 acetone/water mixture. The samples were sonicated in an ice-water bath (10-s x 2) and centrifuged during each washing step. Finally, the sample tubes were heated in a bath at ∼95°C for 2 min to evaporate.

### Immunoblotting and Densitometric Analysis

Immunoblotting was performed using SDS/PAGE on a 15% gel, followed by electroblotting onto a polyvinylidene fluoride (PVDF) membrane, as previously described [33, 37]. The Abs used in the experiments are listed in Supplementary Table S9. Immunoreactivity was detected using horseradish peroxidase (HRP)-conjugated secondary Abs via a chemical fluorescent method as previously described [33, 37]. For the subcellular fractionation assay, after SDS/PAGE and transferred to a PVDF membrane, the membrane was briefly rinsed with water, followed by Tris-buffered saline containing 0.1% Tween-20 (TBT), then wrapped in plastic wrap and cut into three parts, higher than 75 kDa (upper), 75–25 kDa (middle), and below 25 kDa, using a cutter based on the position of the prestained SDS/PAGE standard molecular weight markers. It was sometimes cut into four parts: higher than 75 kDa, 75–40 kDa, 40–20 kDa, and lower than 20 kDa. Each filter was blocked with 0.1 % skim milk in TBT for over 2 h at room temperature and then incubated at 4°C overnight with Abs against Rasa 1 (upper), CFDP1 (middle), and HstH3 (lower). The upper and middle membrane parts were subjected to stripping, followed by blocking, and processed to access by Abs against nestin, NonO for a 75– 40 Da range membrane, and Vdac1 for a 40–25 or 40–20 kDa range membrane. Finally, the middle or 40–20 kDa range membrane was stripped and then incubated with an HRP-conjugated anti-Gapdh Ab after blocking. Multi-box strays were routinely used for all incubation procedures of blocking, reaction with Abs, and washing (Ina Optika MWB-06 or MWB-4). All images were captured at 5.51M Pixel via a GeneGNone-5 (Syngene Bio Imaging) using GeneSnaps software and saved as tiff files. The densitometric evaluation of Western blot signals was conducted using the Band/Peak Quantification plugin in Fiji software [69].

### Mass Spectroscopy Analysis

The Coomassie brilliant blue (CBB)-stained bands or Western blot-positive bands in the ∼43-kDa range in the SDS/PAGE gel were excised and destained, and the samples were processed in duplicate as previously described [10, 33, 37]. They were subjected to trypsin digestion and subsequently analyzed using nano LC-MS/MS with a Q Exactive mass spectrometer. Each sample was run in duplicate or triplicate for mass spectrometry analysis. In the study of the subcellular fractionated samples, SDS/PAGE was performed, and electrophoresis was stopped upon the entry of the samples into the separation gel to treat each as a single band. Each band was excised and subjected to reductive acrylamidation. The samples were digested with trypsin, and equal amounts of each were analyzed in triplicate using the Q Exactive HF-X. The data were analyzed using Proteome Discoverer 3.0 and MASCOT server 2.8 for protein identification and quantification (Supplementary Table S2). For examination of Cyt#1/#2* ratios, each fraction precipitated with TCA was also subjected to brief SDS/PAGE as described above. Each protein concentration was determined based on amino acid composition and analyzed in triplicate. Data were acquired using the Proteome Discoverer 2.4 and MASCOT server 2.7 (Supplementary Table S5). The data of Tables S2 and S5 are available via ProteomeXchange with the identifier PXD063720 http://proteomecentral.proteomexchange.org/. The distribution of Cyt#1 to Cyt#2* ratios was presented as violin plots, and their statistical analysis was performed using EZR [70] (https://cran.r-project.org/web/packages/RcmdrPlugin.EZR/RcmdrPlugin.EZR.pdf).

## Acknowledgments

We thank Dr. Tsutomu Kumazaki for the puromycin-resistant AB2.2 cell line. We appreciate Drs. Shinya Watanabe and Dimitris Beis for information on the properties of Bcnt/cfdp1. We acknowledge Drs. Yan Fei Gao, Gereon Poschmann, Lauri A. Aaltonen, and Justin Cotney for sharing unpublished data. We also thank the following scientists for their valuable insights during this research: Drs. Satoshi Yoshitome, Yasuhiro Kashino, Atsushi Iwai, Masahiro Nishibori, Naoyuki Kataoka, Richard Schulz, Heiko Heerklotz, Song-Tao Liu, Takuto Shimizu, and Mitsuru Okuwaki. We were encouraged by Dr. Stephane Twigg for his comments regarding the nomenclature of the CFDP1 gene. We are willing to thank Editage (www.editing.com) for precision editing the manuscript. We were indebted to Professors Si-Young Song, Hiromi Nochi, and Takashi Tominaga for their support and encouragement.

## Footnote references

^‡^The official name of *Bcnt/Cfdp1* is *Cfdp**1***. However, genetic studies of disease (1) or transcriptome analysis (2) do not provide solid evidence for its involvement in craniofacial development, including tooth development (3). Therefore, the nomenclature of *Cfdp1* is misleading and may need to be more accurate. To maintain consistency with previous studies, the name *Bcnt/Cfdp1* is used instead of *Cfdp**1*** in this article.

## Footnotes

‡ The official name of *Bcnt/Cfdp1* is *Cfdp**1***. However, genetic studies of disease [1] or transcriptome analysis [2] do not provide solid evidence for its involvement in craniofacial development, including mouse tooth development [3]. Therefore, the nomenclature of *Cfdp1* is misleading and needs to be more accurate. To maintain consistency with previous studies, the term *Bcnt/Cfdp1* rather than *Cfdp1* is used in this article.

