## Supplemental Figures 1-4 for "Macromolecular Crowding-Affected Mobility of Bcnt/Cfdp1, a Disordered Component of the Srcap Complex, from the Nucleus to Cytosol in Subcellular Fractionation"

**Supplementary Figure S1**

**No evidence of liquid-liquid phase separation of BCNT/CFDP1 in optoDroplet assay**

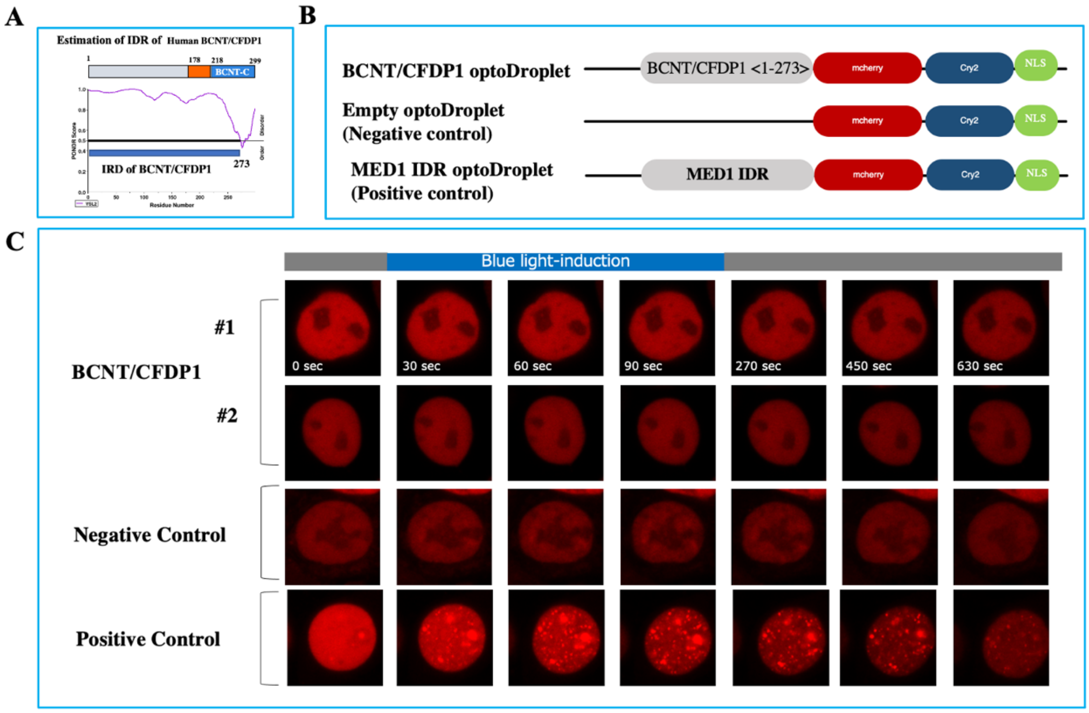


### Legends

**A.** The intrinsically disordered region (IDR) of BCNT/CFDP1 was estimated using the PONDR VSL2 algorithm (http://www.pondr.com/).

**B.** A schematic representation of the constructed plasmids for the optoDroplet assay is presented. BCNT/CFDP1 IDR was amplified by PCR and the obtained fragment was inserted into an optoDroplet vector. The MED1 IDR was used as a positive control, while the empty IDR served as a negative control.

**C.** HEK293T cells were transfected with the three plasmids depicted in B, and the subsequent LLPS activity was measured 48 h later as follows. Blue light stimulation (488 nm stimulus/3 sec, interval/17 sec, mCherry imaging/10 sec, 3 times) was followed by mCherry imaging (imaging/10 seconds, interval/50 seconds) 10 times.

**Supplementary Figure S2**

**Enrichment of the ~43kDa band by Western blot monitoring with antibody**

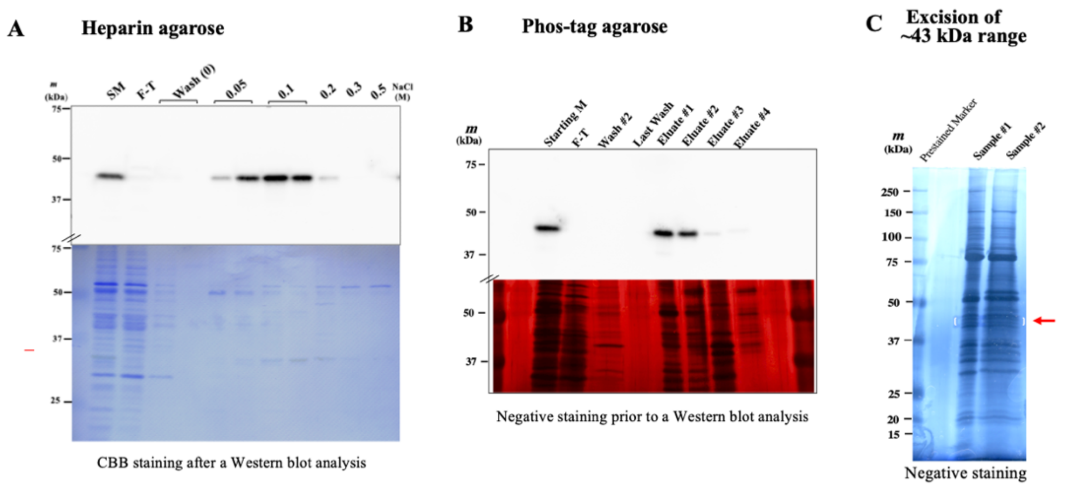


**D**

| Sample | Description | Sum PEP Score | Coverage [%] | # Peptides | # PSMs | # Unique Peptides | Score Mascot: Mascot | # Peptides (by Search Engine): Mascot |
|---|---|---|---|---|---|---|---|---|
| #1 | Bcnt/Cfdp1 | 21.578 | 29 | 7 | 9 | 7 | 180 | 7 |
| #2 | Bcnt/Cfdp1 | 34.399 | 38 | 11 | 14 | 11 | 258 | 11 |

### Legends

The ~43 kDa band was enriched from the cell extracts of the mouse ES AB2.2 cell line using the two sequential columns, heparin agarose **(A)** and Phos-tag resin **(B)**. A constant volume (15 µL) of each fraction was subjected to Western blot analysis. In (A), the filter was stained with CBB after obtaining the image, while the gel was stained with silver staining in (B) prior to Western blotting, shown in the bottom panel, respectively. SM and F-T indicate the starting materials and flow-through fraction, respectively. The eluates from the Phos-tag resin (E1 and E2) were stored in two different buffers. Sample #2 buffer was designed for two-dimensional gel electrophoresis. Following SDS/PAGE separation, the gel was stained using a silver staining kit **(C)**. A gel piece in the range of ~43 kDa (indicated by a red arrow) was excised for mass spectroscopy analysis. The data on Bcnt/Cfdp1 for each sample was extracted from Supplementary Table S1 and is presented in Table D.

### Supplementary Figure S3

**Isolation of FLAG-mouse Bcnt/Cfdp1 using anti-FLAG Ab-conjugated agarose beads**

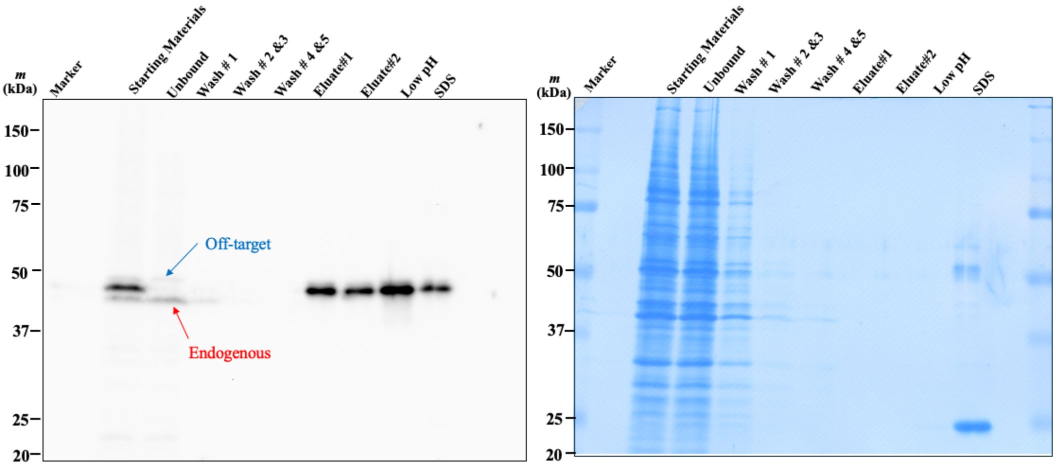


### Legends

C2C12 cells transfected with Flag-tagged mouse *Bcnt/Cfdp1* were cultured nearly confluent, and the cell layers were scraped in RIPA buffer. The supernatant of extracts was incubated with anti-FLAG Ab-conjugated beads. After washing with RIPA buffer, the bound proteins were eluted with DYKDDDDK peptide, glycine-HCl, pH 2.5 buffer (Low pH), followed by boiling in SDS/PAGE sample buffer. The constant volume of each fraction was subjected to Western blot analysis using A305-624-M followed by HRP-conjugated anti-rabbit IgG Ab. Finally, the filter was stained with CBB (right panel).

**Supplementary Figure S4**

**Appearance of an additional 50 kDa band during C2C12 differentiation**

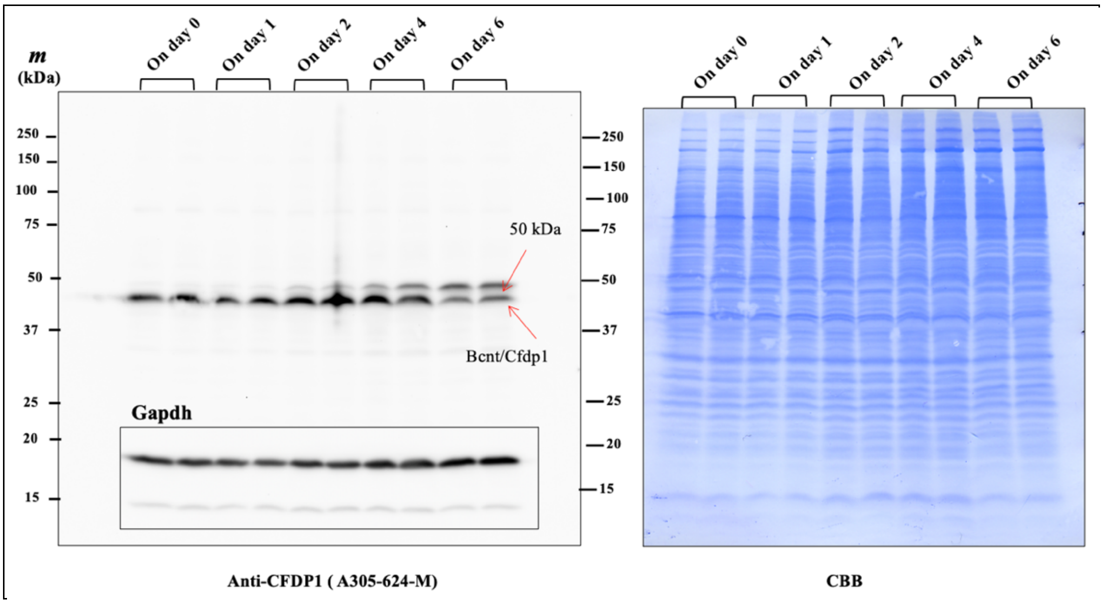


### Legends

C2C12 cells were cultured in duplicate, and the medium was replaced with DMEM-2% horse serum (designated as day 0). On the corresponding day (shown at the top), each cell extract was collected by scraping it in an SDS-containing buffer, boiling it, and then sonicating it. A constant volume of each extract (equivalent to 7.5 μL) was subjected to Western blot analysis after treatment with SDS/PAGE sample buffer. After detecting signals using A305-624-M, the filter was stripped and treated with an anti-HRP-conjugated Gapdh antibody. After obtaining all images, the filter was stained with Coomassie brilliant blue (right panel).
